# Gut Microbiota Dysbiosis-Mediated Gut NLRP3 Inflammasome Activation Exacerbates Corticospinal Tract Injury After Intracerebral Hemorrhage

**DOI:** 10.64898/2026.02.23.707330

**Authors:** Meiqin Zeng, Meichang Peng, Hao Tian, Chen He, Lu Zhang, Wen Yuan, Jiandong Huang, Haitao Sun

## Abstract

Intracerebral hemorrhage (ICH) causes severe neurological deficits, largely attributable to corticospinal tract (CST) injury. However, the underlying mechanism remains unclear, which hinders the development of effective treatment methods. Here, we found that CST injury is not only associated with the activation of NLRP3 inflammasome in the surrounding area of the hematoma as expected, and dysfunction of the blood-brain barrier, but also related to the severe imbalance of the gut microbiota, the activation of NLRP3 inflammasome in the gut and the impairment of gut barrier function after ICH. We therefore systematically investigated how the gut NLRP3 inflammasome and gut dysbiosis exacerbate CST injury after ICH. Knockdown of colon NLRP3 significantly attenuated CST injury and downregulated inflammasome signaling in both the peripheral circulation and the peri-hematomal brain. Consistently, antibiotic-mediated gut microbiota depletion suppressed NLRP3 activation in the gut and brain, improved neurological function, and reduced CST damage. Crucially, fecal microbiota transplantation (FMT) from ICH donors established that the exacerbation of CST injury is dependent on gut dysbiosis. While FMT induced severe pathology in control mice, this effect was improved in gut NLRP3 knockdown recipients, demonstrating that gut NLRP3 is essential for mediating the harmful effects of microbiota dysbiosis. Our findings described a causal gut-brain axis in ICH, wherein microbiota dysbiosis activates the gut NLRP3 inflammasome to exacerbate CST injury, thereby identifying the gut NLRP3 inflammasome as a promising therapeutic target.

## 1. Introduction

Intracerebral hemorrhage (ICH), a devastating stroke subtype caused by bleeding within the brain parenchyma, represents a major public health challenge due to its high disability and fatality rates, as well as its significant economic consequences ^[1]^. Patients often suffer from severe and persistent neurological deficits despite initial recovery, highlighting the limitations of current therapeutic strategies ^[2]^. Moreover, our previous study has shown that this neurological motor dysfunction is attributable not only to peri-hematomal corticospinal tract (CST) injury, but also to distal CST lesions in the cervical spinal cord ^[3]^. Despite these insights, the complete pathological mechanisms driving this widespread CST injury remain incompletely defined, hindering the development of targeted neuroprotective treatments.

In recent years, accumulating evidence has demonstrated that the microbiota-gut-brain axis plays a crucial role in maintaining brain homeostasis and in the pathophysiology of major neurological and psychiatric disorders, such as cognitive impairment, Alzheimer’s disease (AD), multiple sclerosis (MS), stroke, and intracranial aneurysms (IA), through multiple pathways including immune, endocrine, metabolic, and neural mechanisms ^[4]^. Consistent with this, gut microbiota dysbiosis is documented in both clinical patients and experimental models ^[3b, 4e, 5]^. Moreover, preliminary work using pharmacological interventions that ameliorate CST injury in mice concurrently reshape the gut microbial community, suggesting a potential link between post-ICH dysbiosis and neurological injury ^[3b, 6]^. This suggested that targeting the post-ICH dysbiosis could influence the injury outcome, although the underlying mechanisms remain to be elucidated.

Notably, the nucleotide-binding oligomerization domain-like receptor family pyrin domain-containing protein 3 (NLRP3) inflammasome plays a pivotal role in coordinating host physiological functions and regulating central and peripheral immune/inflammatory responses, with its significance being particularly pronounced in neurological disorders ^[4a, 7]^. This complex can sense microbial/danger-associated molecular patterns (MAMPs/DAMPs), triggering caspase-1-dependent IL-1β maturation and propagating inflammation ^[8]^. Recent evidence supports the concept of a microbiota-gut-inflammasome-brain axis, wherein gut-derived NLRP3 activation can propagate systemic inflammation that exacerbates neuropathology ^[4a, 9]^. For instance, fecal microbiota from Alzheimer’s patients can trigger gut NLRP3 activation, accelerating neuroinflammation in recipient mice ^[4b]^. Similarly, in sleep deprivation models, knockdown of gut NLRP3 alleviates neuroinflammation and cognitive impairment ^[8b]^. Notably, our prior work provided the first indication of this axis in ICH, showing that pharmacological NLRP3 inhibition with MCC950 attenuated CST injury and was associated with modifications in the gut microbiota ^[3b]^. While this implied a potential interaction between the gut NLRP3 inflammasome and microbiota in post-ICH pathology, the precise role of gut NLRP3 and its functional dependence on microbiota dysbiosis has not been directly investigated.

Therefore, we hypothesized that ICH-induced gut dysbiosis activates the intestinal NLRP3, which in turn exacerbates CST injury. As the first systematic research of the gut microbiota-NLRP3 axis in post-ICH CST injury, our study establishes that gut NLRP3 activation, driven by ICH-induced dysbiosis, critically aggravates CST pathology. These findings reveal a novel gut-brain communication pathway and nominate the gut NLRP3 inflammasome as a promising therapeutic target for ICH.

## 2. Materials and methods

### 2.1. Animals and study design

We used male C57BL/6J mice (8–10 weeks, 25–30 g) purchased from the Guangdong Medical Laboratory Animal Center and NLRP3 knockout (NLRP3-KO, 8–10 weeks, 25–30 g) mice on a C57BL/6J background obtained from the Jackson Laboratory. Mice were housed in the Experimental Animal Research Center at Southern Medical University. All experimental procedures were approved by the Zhujiang Hospital Ethics Committee (Approval No. LAEC-2022-135) and performed in compliance with the National Institutes of Health (NIH) guidelines for animal care.

The study was conducted through five sequential experiments. In Experiment 1, to establish baseline pathology, mice were randomized into Sham and ICH groups. Experiment 2 investigated the role of NLRP3 by assigning mice to four groups: WT + Sham, NLRP3^-/-^+ Sham, WT + ICH, and NLRP3^-/-^ + ICH. Experiment 3 assessed the function of the gut NLRP3 inflammasome using mice receiving either control AAV-GFP or AAV-mediated NLRP3 knockdown (AAV-NLRP3) before ICH. In Experiment 4, the role of gut microbiota was examined by treating mice with either a vehicle or an antibiotic cocktail (ABX) before ICH induction. Finally, Experiment 5 determined the dependency of gut NLRP3 on the microbiota by colonizing AAV-NLRP3 or AAV-GFP treated mice with ICH microbiota via FMT, alongside an AAV-GFP vehicle control group.

### 2.2. Experimental ICH model and treatments

The collagenase-induced ICH model was established under strict stereotaxic control as previously described with some small modifications ^[3]^. Briefly, mice were anesthetized using 4% isoflurane for induction and maintained at 1.5–2% isoflurane. The mouse was placed in a stereotaxic apparatus (RWD Life Science Co., Shenzhen, China) of a rodent-specific adapter. Next, an incision was made along the midline of the scalp to expose the skull and locate the bregma. Subsequently, a hole was drilled in the right hemisphere of the skull, positioned 2.0 mm lateral to the midline and 0.2 mm anterior to the bregma. Type IV collagenase (Sigma-Aldrich, St. Louis, MI, United States; 0.075 U in 0.5 μL of saline) was infused into the parenchyma at a depth of 3.5 mm over 5 minutes (0.1 μL /min) via a microinjection needle. The needle was held for 10 minutes post-injection to prevent backflow and the slowly withdrawal. Sham controls underwent identical procedures, excluding collagenase administration (0.5μL saline only). The mice were maintained at a temperature of 37.0 ± 0.5 °C throughout the experimental and recovery periods. After the operation, the mice had free access to food and water.

### 2.3. Motor function assessment

We used the open-field test and gait analysis to evaluate the long-term neurologic function of mice following ICH. Our assessments were conducted before ICH and on days 1, 3, 7 and 14 after ICH.

#### 2.3.1. Open-field test

The open-field test was performed as previously described ^[10]^. All mice would be placed in an empty open-field apparatus (40 × 40 × 40 cm) for 5 min, record the movement and distance of the mice. The total distance moved reflects the motor function of the mice. Data were collected for 5□min per mouse and analyzed with EthoVision XT15 software.

#### 2.3.2. Gait analysis

Gait analysis was performed using the Noldus CatWalk XT system (Version 10, Noldus Information Technology). Before the formal tests, mice were trained for 3 days. During the trial, mice were placed at the runway entrance and allowed to traverse freely to the opposite end. A minimum of three uninterrupted runs per session was recorded for each animal to ensure data consistency. Detailed gait parameters, including inter-limb coordination, stride length, and base of support, were automatically quantified by the CatWalk XT software. Critically, the Regularity Index (RI) was calculated as a validated metric of motor coordination deficits, defined as the percentage of normal step sequence patterns relative to total paw placements during locomotion ^[11]^.

### 2.4. Tissue and blood collection

Upon completion of behavioral assessments, mice were deeply anesthetized with 5% isoflurane. Blood was collected into centrifuge tubes, allowed to clot at room temperature for 1 hour, then centrifuged at 3,000 × g for 15 min to isolate serum. Serum was frozen in liquid nitrogen quickly and then stored at -80°C for subsequent cytokine quantification by ELISA. Immediately, after perfusion with ice-cold PBS (pH 7.4), brain and colon tissues were removed. For molecular analyses, brain and colon tissues were frozen in liquid nitrogen and stored at -80°C for protein/RNA extraction required for Western blot (WB) and quantitative PCR (qPCR) experiments. For histology, parallel tissue samples were collected and fixed in 4% paraformaldehyde (PFA) for 48 hours at 4°C, and dehydrated with 20% sucrose for 2 days, followed by dehydration with 30% sucrose for 2 days before immunofluorescence staining.

### 2.5. Immunofluorescence and Image Analyses

For immunofluorescence, sections of the spinal cord (10 µm thick) and brain (15 µm thick) were fixed with methanol for 15 min, penetrated with 0.5% Triton X-100 for 5min and blocked with 5% goat serum for 1 hour at room temperature. Then, these sections were incubated overnight at 4□ with primary antibodies: rat anti-MBP (1:400; Novus, NB600-717), mouse anti-SMI-32 (1:200; BioLegend, 801701), rabbit anti-NF200 (1:400; Sigma, N4142), rat anti-GFAP (1:400; Invitrogen, 13-0300), and rabbit anti-Iba1 (1:400; Abcam, Ab178846). After PBS washes, slices were then incubated with Alexa Fluor-conjugated species-matched secondary antibodies (1:400; Abcam, Ab150077 and Ab150160) for 1 hour at room temperature. Then, the slices were sealed by anti-quench sealant DAPI and captured by a Nikon Eclipse Ti2 microscope, and immunofluorescence intensity and the number of positive cells were analyzed using ImageJ.

### 2.6. Total protein extraction and western blot

Twenty milligrams of brain or colon tissues were homogenized in cold RIPA lysis buffer (Beyotime, P0013B) containing a 1:100 protease inhibitor mixture (Beyotime, P1005), and then two grinding beads were used for each sample. Then, samples were centrifuged at 14,000 × g at 4 ° C for 15 minutes. The supernatant was collected, and the protein concentration was detected using the BCA assay kit (Beyotime, P0010S). Next, the loading buffer (Fudebio, FD002) was added to the supernatant in a 1:4 ratio and boiled at 95°C for 15 minutes, after which it was stored at -80°C. An equal amount of total protein (30-50μg) was separated on SDS-PAGE (Bio-Rad) gels and then transferred to 0.22μm polyvinylidene fluoride (PVDF) membranes (Millipore, ISEQ00010). The membranes were blocked at room temperature in 5% non-fat milk for 1 hour, then incubated overnight at 4 □ with following primary antibodies: rabbit anti-NLRP3 (1:1000, immunoway, YM8024), rabbit anti-cleaved Caspase-1 p20 (1:1000, abmart, P79884R2M), rabbit anti-IL-1β (1:1000, MCE, HY-P80720), rabbit anti-ZO-1 (1:5000, proteintech, 21773-1-AP), rabbit anti-Occludin (1:5000, proteintech, 27260-1-AP), rabbit anti-β-tubulin (1:5000, affinity, AF7011), and rabbit anti-actin (1:5000, Beyotime, AF5006). After primary antibody incubation, membranes were washed 3 times in TBST for 10 minutes and then incubated with secondary antibody at room temperature for 1 hour. Signals were detected using FDbio-Dura ECL substrate (Fudebio, FD8020) and captured on a UVITEC Alliance Micro Q9. Images were quantitatively analyzed using ImageJ software.

### 2.7. Total RNA extraction and Quantitative Real-time RT-PCR

Total RNA was isolated from peri-hematomal brain and colon tissues using TRIZOL reagent (Invitrogen, 15596026) following the manufacturer’s protocol. Briefly, tissues were homogenized in 1 mL TRIZOL with two grinding beads, then using a cryogenic grinding machine at 70 Hz for 4 min total (8 cycles of 30 s homogenization/10 s pause, 4°C). After complete lysis, 200μL chloroform was added, samples were vortexed vigorously for 30s, then centrifuged at 12,000 rpm for 15min at 4°C. The aqueous phase (450μL) was transferred to new tubes, mixed with an equal volume of isopropanol, and incubated at 4°C for 30 min. RNA pellets were obtained by centrifugation (12,000 rpm, 10min, 4°C), washed twice with 80% ethanol 7,500 rpm, 5min), air-dried for 5min, and resuspended in 30μL nuclease-free water (Beyotime, R0022). cDNA was synthesized from 1μg total RNA using Evo M-MLV RT MIX Kit (Accurate Biology, AG11728) under standard conditions: 37°C for 15 min → 85°C for 5s. qPCR reactions employed SYBR Green Pro Taq HS qPCR Kit (Accurate Biology, AG11701) on QuantStudio 3 Pro system (Applied Biosystems) with cycling parameters: stage 1 (hold stage, 95°C for 30s); stage 1(40 cycles of 95°C for 5s → 60°C for 30s); stage 3 (Melt curve: 95□ for 15s→55□ for 1min→95□ for 1s). Data were normalized to *Actb* reference genes and analyzed by the 2^(-ΔΔCt) method. The specific sequences of primers used are shown in Table S1.

### 2.8. Antibiotic treatment

ABX (antibiotic cocktail) represents depleting the gut microbiota in mice using a combination of broad-spectrum antibiotics including (0.5 g/L, 94747; Sigma-Aldrich), ampicillin (1 g/L, A9518; Sigma-Aldrich, St. Louis, MO, USA), neomycin (1 g/L, N1876; Sigma-Aldrich), and metronidazole (1 g/L, M3761; Sigma-Aldrich) adapted from established protocols ^[12]^. And this method has already been confirmed can successfully clear the gut microbiota ^[13]^. These antibiotics were dissolved in the sterile drinking water provided to the mice daily. Considering metronidazole palatability issues, a gradient adaptation protocol was implemented: no metronidazole on day 0, 0.25g/L concentration from day 2, 0.5g/L from day 6, and reaching the full concentration of 1g/L from day 9. Mice received ABX treatment for 14 days before ICH induction, with continuous administration until sacrifice. Fresh antibiotic solutions were replaced every 72 hours to maintain potency. Control groups received autoclaved water.

### 2.9. Fecal microbiota transplantation (FMT)

Fecal microbiota transplantation (FMT) was performed via oral gavage as previously described, with minor modifications ^[14]^. Mice first underwent a 2-week antibiotic pretreatment to clear the gut microbiota, followed by a 48-hour recovery period without antibiotics. Subsequently, recipients received daily FMT for 7 consecutive days before ICH induction. FMT administration was continued after ICH modeling until the end of the experiment. Briefly, fresh fecal samples were collected from ICH donor mice immediately after defecation. Fecal pellets from multiple donors were pooled and homogenized in anaerobic PBS (200 mg per 2 mL) by vortexing for 5 minutes. The homogenate was then gently centrifuged at 1200 × g for 3 minutes to sediment large particulate matter. Recipient mice received 200 μL of the final suspension daily via oral gavage.

### 2.10. NLRP3 knockdown by an AAV delivery approach

For knockdown of colon NLRP3, AAV-U6-shRNA (nlrp3)-CMV-EGFP-pA and its control AAV-U6-shRNA (scramble)-CMV-EGFP-pA were delivered into the colon, which were constructed and purchased from BrainVT A (Wuhan) Co., Ltd. The sequence used for RNAi targeting NLRP3 was GGAGGACAGCCTTGAAGAAGA, while the non-specific sequence (CCTAAGGTTAAGTCGCCCTCG) served as a negative control. Vectors encoding enhanced green fluorescent protein (GFP) were utilized to confirm successful delivery in mice. For colonic enema delivery, adeno-associated virus (AAV) vectors carrying either the NLRP3 gene (AAV-NLRP3) or GFP (AAV-GFP) were administered. Specifically, mice were fasted overnight to allow bowel clearance. Two sequential enemas of 20 mM N-acetylcysteine (NAC, 300 μL; Sigma, A9165) at 15-min intervals. Subsequently, 200 μL AAV suspension (total dose: 5 × 10¹² viral genomes [vg] in saline) was infused into the gut via enema. After administration, mice were maintained in a vertical (head-down) position immediately for 2 min to ensure optimal viral transduction. ICH model induction would happen 3 weeks later. Finally, the effects of in vivo transfection were observed via fluorescence microscopy, qPCR and Western blotting.

### 2.11. Elisa

Peripheral blood samples were centrifuged, and serum (the supernatant) was promptly separated for subsequent analysis. Serum Caspase-1 (MEIMIAN, MM-0820M1-11c) and interleukin-1β (IL-1β, Elabscience, E-EL-M0037) levels were quantified using a commercial enzyme-linked immunosorbent assay (ELISA) kit according to the manufacturer’s instructions.

### 2.12. Fecal sample collection and 16S rRNA gene sequencing analysis

Upon the completion of the ICH model, mice were closely observed until four fresh fecal pellets were collected from each mouse with individual 1.5 ml centrifuge tubes, and stored at -80 □ for microbiome analysis. The microbiota analysis was conducted following the previously described method of our published works ^[3b, 6a, 15]^.

### 2.13. Statistical analyses

All statistical analyses were performed using GraphPad Prism 8.3(GraphPad Software, San Diego, CA). Continuous variables are presented as mean ± standard error of the mean (SEM). Normality and homogeneity of variances were assessed by the Shapiro-Wilk and Levene’s tests, respectively. For data satisfying both assumptions, parametric tests were applied: unpaired Student’s t test for two individual group comparisons and one-way ANOVA with Bonferroni post hoc test for multiple groups (>2 groups). If the data met the assumption of normality but not the assumption of homogeneity of variances, Welch’s corrected t-test and Welch’s ANOVA were applied. If both assumptions were violated, non-parametric tests (e.g., Mann-Whitney *U* test and Kruskal-Wallis test) were used with Dunn’s post hoc correction for multiple groups (>2 groups). Two-way ANOVA with Sidak’s multiple comparisons test used to compare differences between multiple groups occurring over time. Graphical presentations were generated using GraphPad Prism 8.3 and ImageJ. Statistical significance was defined as *p< 0.05. Significance levels are denoted: *p < 0.05, **p< 0.01, ***p < 0.001.

## 3. Results

### 3.1. The Cervical Spinal Cord was the Predominant Site of Distal Corticospinal Tract Injury after ICH

Although people have fully recognized the secondary injury suffered by the corticospinal tract after ICH, the existing research mainly focuses on the area around the hematoma ^[16]^. Our previous work highlighted that distal CST injury, particularly Wallerian degeneration persisting for weeks in the cervical spinal cord, is closely related to long-term motor deficits^[3a]^. However, the impact of this pathological process on other distal spinal cord regions remained unexplored. To explain why we chose the cervical area as the key point for observing distal CST injury in our subsequent experiments, we investigated the ends of the spinal cord, including cervical and lumbar segments, in a mouse ICH model induced by collagenase IV (Fig.1A, B). Interestingly, immunofluorescence analysis revealed significant axonal injury, as indicated by decreased expression of myelin basic protein (MBP) and neurofilament-200 (NF200), localized predominantly to the cervical spinal cord following ICH (MBP: t_6_ = 3.201, p = 0.0186; NF200: t_6_ = 4.052, p = 0.0067, Fig.1C, F). In contrast, the lumbar enlargement showed no comparable downregulation (MBP: t_7_ = 0.9790, p = 0.3602; NF200: t_7_ = 0.9954, p = 0.3527 Fig.1I, L). Furthermore, assessment of neuroinflammation at 14 days post-ICH, using Iba-1 and GFAP as markers for microglia and astrocytes, indicated significantly increased activation within the cervical spinal cord (Iba-1: t_6_ = 2.795, p = 0.0314; GFAP: t_6_ = 4.760, p = 0.0031, Fig.1D-E and G-H). In contrast, the lumbar enlargement showed minimal inflammatory changes (Iba-1: t_6_ = 0.2582, p = 0.8049; GFAP: t_7_ = 0.1145, p = 0.9121, Fig.1J-K and M-N). Collectively, these data identify the cervical spinal cord as the primary locus of distal axonal injury and sustained neuroinflammation after ICH. This established a critical anatomical and pathological foundation for investigating the mechanisms underlying distal CST damage and its contribution to long-term functional impairment.

**Fig. 1.**
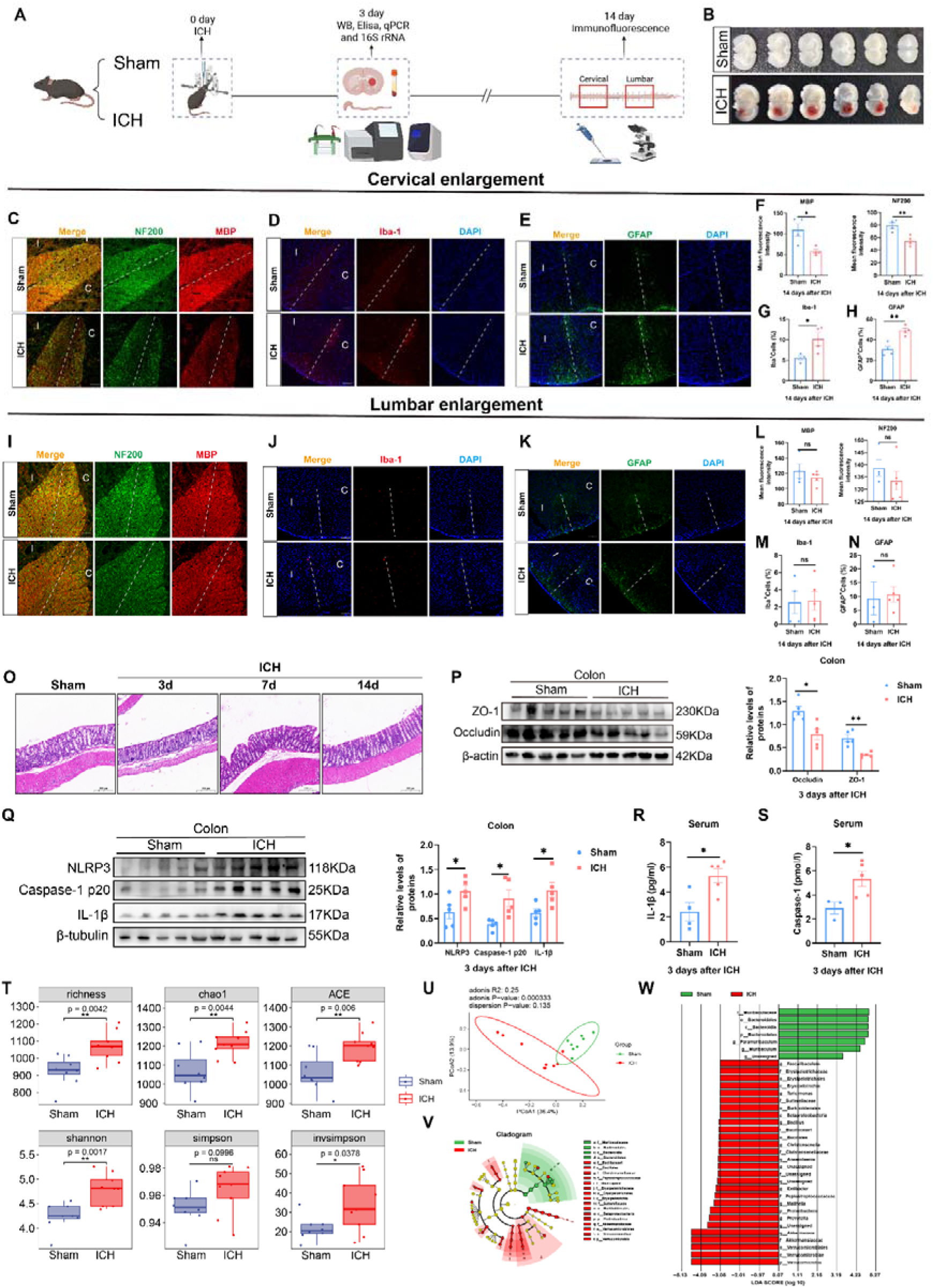
Gut NLRP3 inflammasome activation, gut barrier disruption, microbiota dysbiosis and cervical CST injury following ICH. **(A)** Experimental timeline and methods. Created with BioRender. **(B)** ICH model validation. **(C, F)** Representative immunofluorescence images showing MBP and NF200 expression in the cervical spinal cord (n=4/group). Scale bar: 50 μm. Statistical analysis by the two-tailed t test. **(D-E, G-H)** Representative images of Iba-1 and GFAP immunostaining in the cervical spinal cord at 14 days post-ICH (n=4/group). Scale bar: 100 μm. Statistical analysis by the two-tailed t test. **(I, L)** Representative immunofluorescence images showing MBP and NF200 expression in the lumbar spinal cord (n=4-5/group). Scale bar: 50 μm. Statistical analysis by the two-tailed t test. **(J-K, M-N)** Representative images of Iba-1 and GFAP immunostaining in the lumbar spinal cord at 14 days post-ICH (n=3-5/group). Scale bar: 100 μm. Statistical analysis by the two-tailed t test. **(O)**Representative H&E-stained sections of colon tissue at 3, 7 and 14 days post-ICH. Scale bar: 200 μm. **(P)** Western blot analysis of tight junction proteins (Occludin, ZO-1) in colon tissue. β-actin served as the loading control, n=5/group. Statistical analysis by the two-tailed t test. **(Q)** Western blot analysis of NLRP3, Caspase-1 p20, and IL-1β in colon tissue. β-Tubulin served as the loading control, n=5/group. Statistical analysis by two-tailed t test, except for the Caspase-1 p20 protein data, was analyzed by Welch’s t test. **(R-S)** ELISA measuring levels of Caspase-1 and IL-1β from peripheral blood, n=3-5/group. Statistical analysis by the two-tailed t test. **(T)**Assessment of microbiota α-diversity, n=8-9/group. The InvSimpson index was compared using a two-tailed Welch’s t-test, while other α-diversity indices were compared using a two-tailed Student’s t-test. (U)Principal Coordinate Analysis (PCoA) based on unweighted Bray-Curtis Distance, n=8-9/group (permutational multivariate analysis of variance). **(V-W)** Linear discriminant analysis Effect Size (LEfSe) identifying differentially abundant bacterial taxa, n=8-9/group (Kruskal-Wallis test). Data are presented as mean ± SEM. Significance levels are defined as *p < 0.05, **p < 0.01. C, contralateral; I, ipsilateral; ns, no significance.

### 3.2. ICH Triggered Gut NLRP3 Inflammasome Activation, Barrier Disruption, and Microbiota Dysbiosis

Consistent with prior reports demonstrating NLRP3 inflammasome activation and blood-brain barrier (BBB) disruption following ICH, our study confirmed activation of the NLRP3 inflammasome in the ipsilateral brain hemisphere at 3 days post-ICH ^[17]^. This was evidenced by significantly increased protein levels of NLRP3, IL-1β, and cleaved Caspase-1 p20, alongside the reduction in the BBB tight junction proteins including ZO-1 and Occludin after ICH (NLRP3: t_8_ = 3.943, p = 0.0043; Caspase-1 p20: t_8_ = 5.734, p = 0.0004; IL-1β : t_8_ = 2.376, p = 0.0449; ZO-1: t_8_ = 2.704, p = 0.0269; Occludin: t_8_ = 2.687, p = 0.0276, Fig.S1B, C). Meanwhile, we would like to investigate whether there are any pathological changes in the intestines. Histological examination of colon tissue revealed substantial intestinal barrier injury at 3 days post-ICH, with evidence of residual damage persisting at 14 days (Fig.1O). Based on these findings, the colon of ICH group exhibited NLRP3 inflammasome activation, demonstrated by increased protein levels of NLRP3, IL-1β, and cleaved Caspase-1 p20, concurrent with a decrease in ZO-1 and Occludin (NLRP3: t_8_ = 2.384, p = 0.0443; Caspase-1 p20: Welch-corrected t_4.641_ = 2.995, p = 0.0332; IL-1β : t_8_ = 2.647, p = 0.0294; ZO-1: t_8_ = 4.872, p = 0.0012; Occludin: t_8_ = 3.354, p = 0.01, Fig.1P, Q). These findings were further supported at the mRNA level, as qPCR analysis showed significant upregulation of *Nlrp3*, *Asc*, and *Casp1* (encoding Caspase-1) and downregulation of *Tjp1* (encoding ZO-1) and *Ocln* (encoding Occludin) in colon tissue (*Nlrp3*: Welch-corrected t_4.615_ = 7.175, p = 0.0011; *Casp1*: t_8_ = 5.941, p = 0.0003; *Asc* : t_6_ = 3.632, p = 0.0109; *Tjp1*: Welch-corrected t_4.750_ = 5.304, p = 0.0037; *Ocln*: Welch-corrected t_4.040_ = 3.831, p = 0.0183, Fig.S1A). The systemic inflammatory response was also indicated by elevated concentrations of Caspase-1 and IL-1β in peripheral blood (Caspase-1: t_6_ = 2.633, p = 0.0389; IL-1β: t_7_ =3.106, p = 0.0172, Fig.1R, S).

Given the documented crosstalk between the gut NLRP3 inflammasome and the microbiota in neurological disorders, we profiled the gut microbiota 3 days after ICH ^[9]^. Alpha-diversity was significantly increased across several indices (Richness index, CHAO1 index, ACE index, and Shannon index), while Beta-diversity revealed a clear separation between the ICH and Sham groups (Richness: t_15_= 3.372, p = 0.0042; Chao1: t_15_= 3.351, p = 0.0044; Shannon: t_15_= 3.802, p = 0.0017; ACE: t_15_= 3.200, p = 0.006; Simpson: t_15_= 1.755, p = 0.0996; Invsimpson: Welch-corrected t_11.11_ = 2.356, p = 0.0378, Fig.1T, U). Linear discriminant analysis Effect Size (LEfSe) analysis identified that specific genera such as *Akkermansia*, *Prevotella* and *Extibacter* were enriched in the ICH group, whereas the Sham group was predominantly colonized by *Paramuribaculum* and *Muribaculum* (Fig.1V, W). Stacked bar charts illustrated compositional shifts at phylum, family, and genus levels (Fig.S1D–F). Our research results show that ICH not only activates the NLRP3 inflammasome in the colon but also damages the BBB and intestinal barrier, accompanied by changes in the gut microbiota.

### 3.3. Colon NLRP3 knockdown mitigated neurological deficits and CST injury after ICH

Previous work from our group demonstrated that pharmacological NLRP3 inhibition with MCC950 attenuates peri-hematomal and cervical spinal cord CST injury post-ICH ^[3b]^. To initially validate the therapeutic potential of systemic NLRP3 inhibition, we utilized NLRP3 knockout (KO) mice (NLRP3: t_6_ = 2.806, p = 0.0309; IL-1β : t_6_ = 3.302, p = 0.0164; ZO-1: t_6_ = 16.36, p <0.0001; Occludin: t_6_ = 7.260, p = 0.0003, Fig.S2A, B). ICH mice lacking NLRP3 exhibited improved motor performance (pre: p = 0.9998; 1d: p = 0.1419; 3d: p = 0.0318; 7d: p = 0.0016 Fig.S2 D, E), attenuated peri-hematomal (MBP: F _(3, 13)_ = 8.969, p= 0.0018; NF200: F _(3, 13)_ = 6.199, p= 0.0076; SMI-32: F _(3, 10)_ = 23.46, p<0.0001) and cervical spinal CST (MBP: F _(3, 6.109)_ = 94.40, p<0.0001; NF200: F _(3, 14)_ = 6.571, p= 0.0053) demyelination and neuroinflammation (Iba-1: F _(3, 11)_ = 8.313, p= 0.0036; GFAP: F _(3, 12)_ = 6.583, p= 0.007), confirming NLRP3 as a promising target for intervention (Fig.S2 F-Q).

To further investigate the specific impact of the gut NLRP3 inflammasome on CST injury following ICH, we knocked down NLRP3 expression in the colon using adeno-associated virus (AAV) delivering shRNA targeting NLRP3 (AAV-NLRP3) or control GFP (AAV-GFP) (Fig.2A). Immunofluorescence confirmed successful viral transduction, and qPCR and western blot analyses verified the effective reduction of NLRP3 mRNA and protein in colon tissue, respectively (NLRP3: t_6_ = 8.835, p = 0.0001; *Nlrp3*: t_4_ = 3.696, p = 0.0209, Fig.2B-C and Fig.S3A-B). This knockdown consequently suppressed the activation of the inflammasome pathway, as evidenced by decreased levels of cleaved Caspase-1 p20 and IL-1β (Caspase-1 p20: t_6_ = 3.477, p = 0.0132; IL-1β : t_6_ = 2.510, p = 0.0459; *Casp1*: t_5_ = 3.291, p = 0.0217 Fig.2B-C and Fig.S3A-B). We then assessed the functional and structural consequences of colon NLRP3 knockdown. In behavioral tests, the AAV-NLRP3 group showed significant improvements in motor function. The total distance traveled in the open-field test was increased at days 3, 7, and 14 post-ICH, and CatWalk XT™ gait analysis revealed a higher regularity index compared to AAV-GFP controls (Distance travelled: pre: p= 0.8409; 1d: p= 0.0896, 3d: p=0.0022; 7d: p= 0.1313; 14d: p= 0.0405; Regularity index: pre: p= 0.9542; 3d: p=0.4406; 7d: p= 0.4303; 14d: p= 0.027, Fig.2D-G). Structurally, immunofluorescence analysis of the peri-hematomal region demonstrated that colon NLRP3 knockdown mitigated the ICH-induced loss of MBP (MBP: t_4_ = 3.034, p = 0.0386; NF200: t_4_ = 3.124, p = 0.0354, Fig.2H, J, K). SMI-32/MBP co-staining further confirmed a significant reduction in the proportion of degenerating axons within the peri-hematomal CST (SMI-32/MBP: t_5_ = 2.998, p = 0.0302, Fig.2I, L). Critically, these protective effects were also observed in the distal cervical spinal cord. At 14 days post-ICH, colon NLRP3 knockdown restored the immunofluorescence intensity of the axonal markers MBP and NF200 in the cervical CST (MBP: t_5_ =3.528, p = 0.0168; NF200: t_5_ = 6.793, p = 0.0011, Fig.2M, P, Q), and concurrently suppressed neuroinflammation, as indicated by reduced expression of Iba-1 (microglia) and GFAP (astrocytes) (Iba-1: Mann-Whitney U = 0, p = 0.0286; GFAP: t_5_ = 3.119, p = 0.0263, Fig.2N, O, R, S).

**Fig 2.**
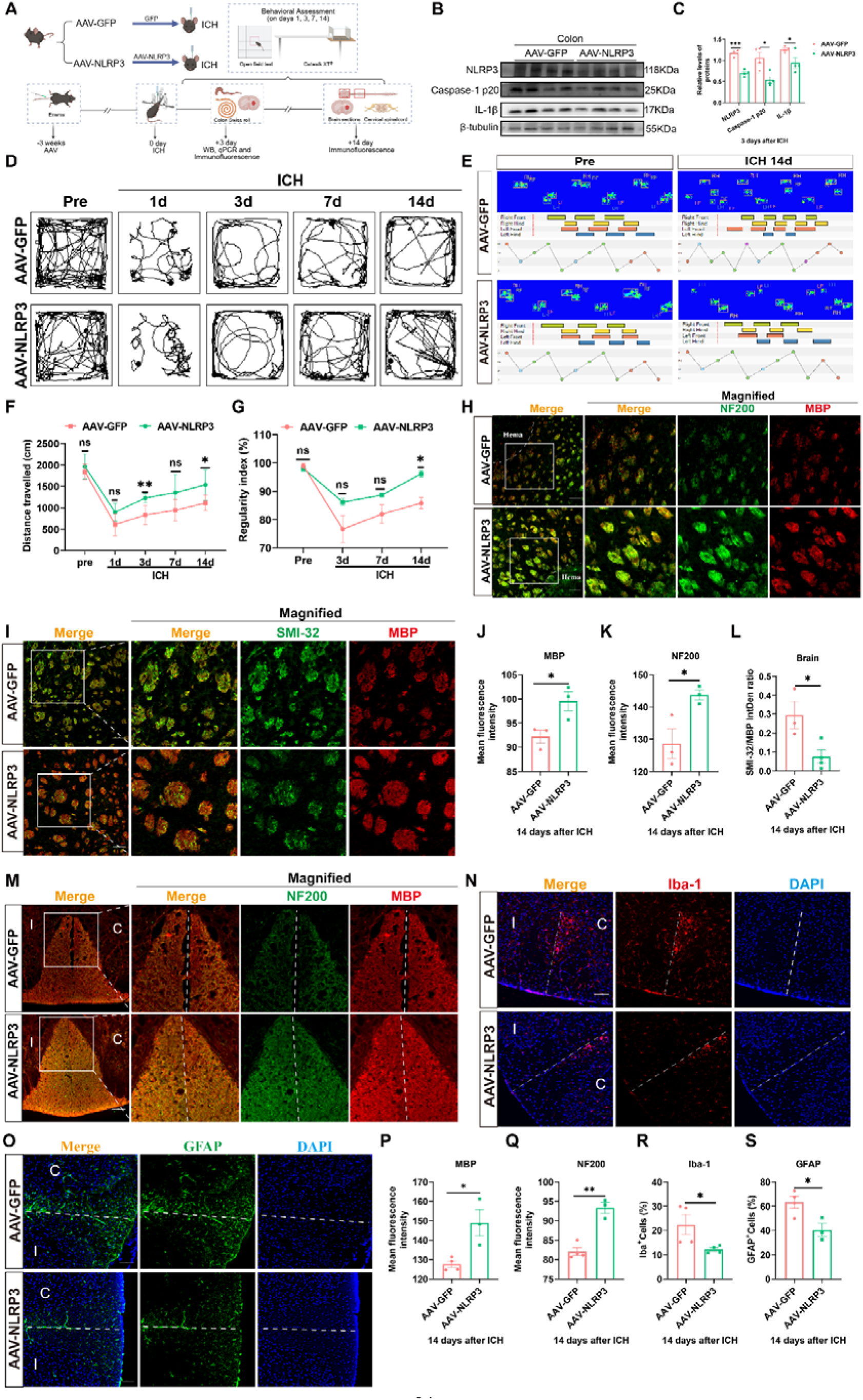
Colon NLRP3 knockdown ameliorated neurological deficits and attenuates CST injury after ICH. **(A)** Schematic diagram of the experimental timeline and treatment groups, created with BioRender (https://BioRender.com/s5rgq47). **(B, C)** Western blot analysis of NLRP3 inflammasome components (NLRP3, Caspase-1 p20, IL-1β) in colon tissues from the indicated groups, n=4/group. Statistical analysis by the two-tailed t test. **(D, F)** Representative open-field test images and quantification of total travel distance in the open-field test across timepoints at 1, 3, 7, 14 days (n=9 mice/group). Statistical analysis by two-way ANOVA and Sidak’s multiple comparisons test. **(E, G)** Representative paw print patterns from the CatWalk XT gait analysis system, and the regularity index from each group at 3, 7 and 14 days. n=4 mice/group. Statistical analysis by two-way ANOVA and Sidak’s multiple comparisons test. **(H, J, K)** Representative MBP and NF200 immunofluorescence staining in peri-hematoma brain sections. n=3 mice/group. Scale bar: 100μm. Statistical analysis by the two-tailed t test. **(I, L)** Representative MBP and SMI-32 immunofluorescence staining in peri-hematoma brain sections. n=3-4 mice/group. Scale bar: 100 μm. Statistical analysis by the two-tailed t test. **(M, P, Q)** Representative immunofluorescence images and quantification of MBP and NF200 in the cervical spinal cord (n=3-4 mice/group). Scale bar: 100 μm. Statistical analysis by the two-tailed t test. (**N, O, R, S)** Representative images and quantification of Iba-1 (microglia) and GFAP (astrocytes) immunofluorescence in the cervical spinal cord (n=3-4 mice/group). Scale bar: 100 μm. Statistical analysis by the Mann-Whitney test and the two-tailed t test, respectively. Data are presented as mean ± SEM. Significance levels are defined as *p < 0.05, **p < 0.01, ***p < 0.001, and ****p < 0.0001.

To sum up, colon NLRP3 knockdown exhibited the neuroprotective phenotypes observed in global NLRP3-KO mice, ameliorating motor deficits and preserving CST integrity in both peri-hematoma and cervical spinal cord. These results establish the gut NLRP3 inflammasome as a pivotal mediator of post-ICH neural injury and a compelling therapeutic target.

### 3.4. Colon NLRP3 knockdown attenuated systemic and cerebral inflammasome activation and BBB disruption after ICH

To elucidate the mechanisms through which gut NLRP3 knockdown confers neuroprotection, we assessed its impact on inflammasome signaling in the brain and peripheral circulation. Before this, we also discovered that knocking down the gut NLRP3 inflammasome also led to a reduction in the damage to intestinal barrier proteins *Tjp1* and *Ocln* after ICH (*Tjp1*: t_5_ = 4.876, p = 0.0046; *Ocln*: t_6_ = 2.804, p = 0.031, Fig. S2C). Western blot analysis of the ipsilateral hemisphere revealed that colon NLRP3 knockdown significantly suppressed the activation of the NLRP3 inflammasome in the brain, as demonstrated by reduced protein levels of NLRP3, cleaved Caspase-1 p20 and IL-1β compared to AAV-GFP controls (NLRP3: t_6_ =4.562, p = 0.0038; IL-1β : t_6_ = 4.154, p = 0.006; Caspase-1 p20: t_6_ = 2.576, p=0.042; Fig.3A-B).

**Fig 3.**
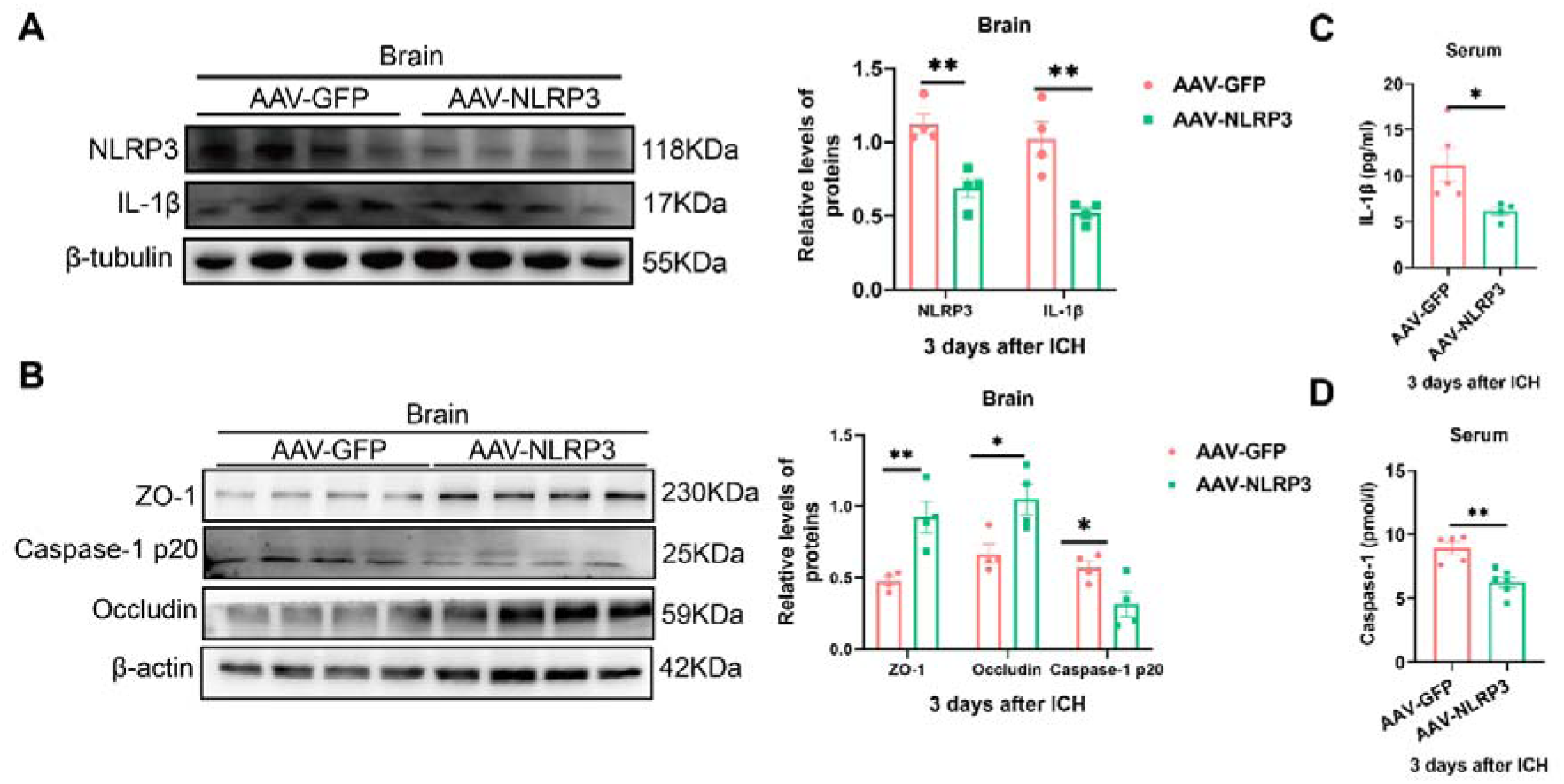
Colon NLRP3 knockdown attenuated systemic and cerebral inflammasome activation and BBB disruption after ICH. **(A-B)** Western blot analysis of NLRP3 inflammasome components (NLRP3, Caspase-1 p20, IL-1β) and tight junction proteins (ZO-1, Occludin) in brain tissue. β-Tubulin was used as the loading control (n=4 mice/group). Statistical analysis by the two-tailed t test. **(C-D)** ELISA quantification of serum levels of IL-1β (C) and Caspase-1 (D) from peripheral blood (n=4-5 and 5-6 mice/group, respectively). Statistical analysis by two-tailed t test with Welch’s correction and two-tailed t test, respectively. All data are presented as mean□±□SEM. Data were considered significant if *p < 0.05, **p < 0.01, ***p < 0.001, ****p < 0.0001.

Given the established link between inflammasome activation and BBB integrity, we further investigated the effect of colon NLRP3 knockdown on this critical structure. We found that the knockdown attenuated BBB disruption, as evidenced by a partial restoration of the tight junction proteins ZO-1 and Occludin in the peri-hematomal region (ZO-1: t_6_ =3.974, p = 0.0073; Occludin : t_6_ = 3.025, p = 0.0232, Fig.3B).

We next evaluated whether this local suppression in the brain was part of a broader systemic effect. Consistent with the cerebral findings, ELISA measurements showed significantly lower concentrations of Caspase-1 and IL-1β in the peripheral blood of AAV-NLRP3 mice at 3 days post-ICH (Caspase-1: t_9_ =4.296, p = 0.002; IL-1β: Welch-corrected t_4.592_ =2.710, p=0.0462, Fig.3C-D).

In summary, colon NLRP3 knockdown mitigated neurological deficits and CST injury, reduced neuroinflammation in the cervical spinal cord, suppressed NLRP3 inflammasome activation in the brain and the systemic circulation, as well as preserved BBB integrity. These findings strongly indicated the gut NLRP3 inflammasome as an important driver of gut-brain axis signaling and multi-compartment pathology following ICH.

### 3.5. Gut microbiota depletion attenuated NLRP3 inflammasome activation in the colon, peripheral blood, and brain after ICH

Having established that gut NLRP3 knockdown alleviates CST injury and that ICH induces significant gut microbiota dysbiosis, we next investigated whether these two phenomena are functionally linked. We hypothesized that gut microbiota dysbiosis is a prerequisite for NLRP3 inflammasome activation after ICH. To test this, we depleted the gut microbiota using a broad-spectrum ABX before ICH induction and assessed outcomes at 3 days post-ICH (Fig.4A). Successful microbiota depletion was confirmed by both cecal enlargement and a marked reduction in bacterial DNA concentration (colon: F_(2, 12)_ = 3.536, Sham vs. ICH+Vehicle, p= 0.9963, Sham vs. ABX+ICH, p= 0.1215, ICH+Vehicle vs. ABX+ICH, p= 0.0663; cecum: F (2, 12) = 82.26, Sham vs. ICH+Vehicle, p= 0.6859, Sham vs. ABX+ICH, p <0.0001, ICH+Vehicle vs. ABX+ICH, p <0.0001; Bacterial load: t_5_=13.04, p<0.0001 Fig.S4A, C).

**Fig 4.**
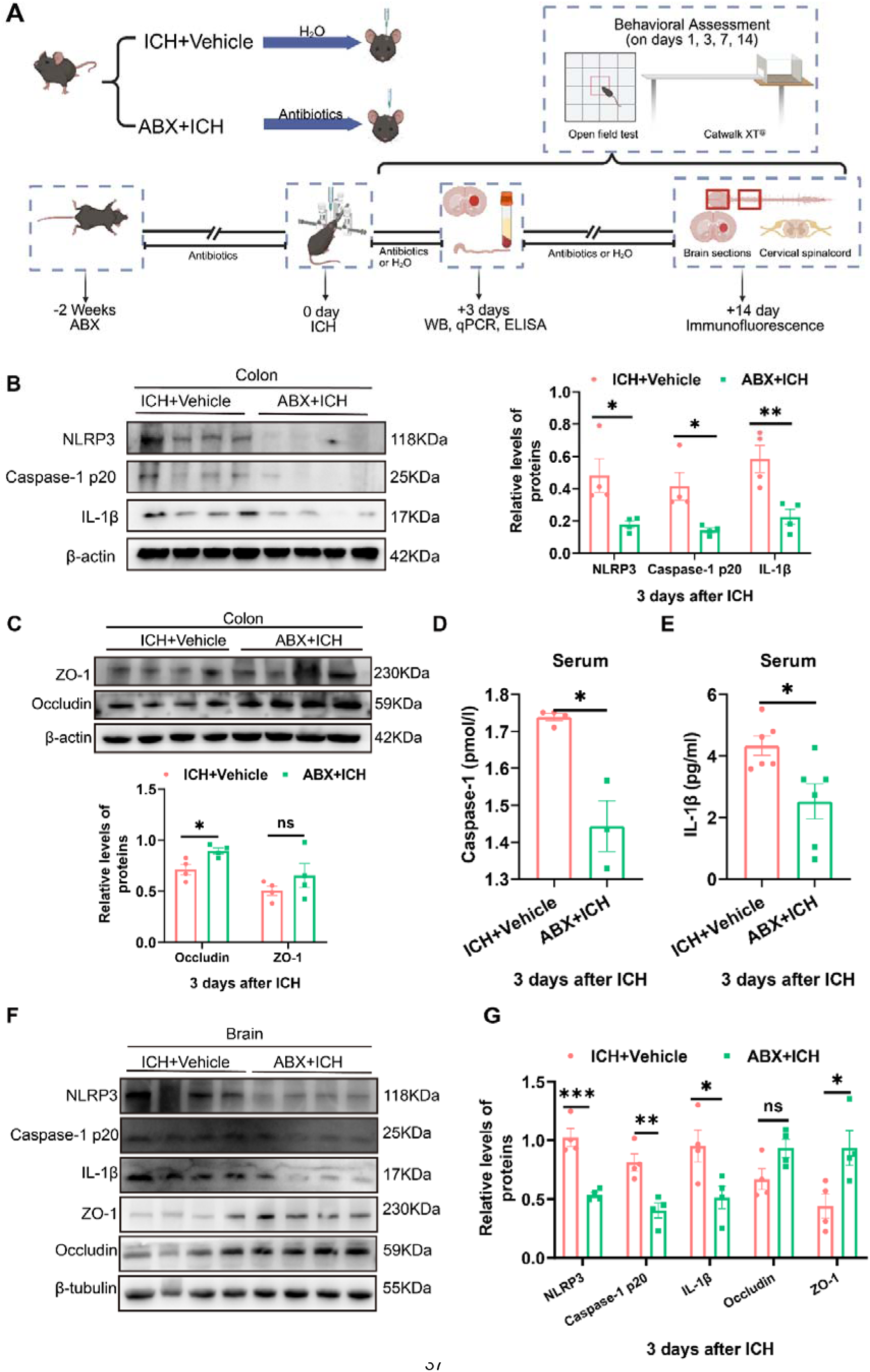
Gut microbiota depletion attenuated NLRP3 inflammasome activation in colon, peripheral blood, and brain after ICH. **(A)** Summary of the experimental procedures, timepoints, created by BioRender (https://BioRender.com/dmwrlny). **(B-C)** Western blot analysis of NLRP3 inflammasome components (NLRP3, Caspase-1 p20, IL-1β) and tight junction proteins (ZO-1, Occludin) in colon tissue (n=4 mice/group). β-Actin was used as the loading control. Statistical analysis by the Mann-Whitney test, except for the IL-1β protein data, was analyzed by the two-tailed t test. **(D)** Elisa analysis of serum Caspase-1 from peripheral blood. n=3-4 mice/group. Statistical analysis by two-tailed t test with Welch’s correction**. (E)** Elisa analysis of serum IL-1β from peripheral blood. n=6 mice/group. **(F-G)** Western blot analysis of NLRP3 inflammasome components (NLRP3, Caspase-1 p20, IL-1β) and BBB proteins (ZO-1, Occludin) in brain tissue in brain tissue. n=4 mice/group. β-tubulin was used as the loading control in Western blot analysis. Statistical analysis by two-tailed t test. Data are presented as mean ± SEM. Significance was set at *p < 0.05, **p < 0.01, ***p < 0.001, and ****p < 0.0001.

To ensure that any subsequent group differences were not attributable to pre-existing variations in baseline health, we confirmed that all mice had comparable body weight (body weight: pre: p = 0.477; 3d: p = 0.0195; 5d: p = 0.0009; 7d: p = 0.023; 9d: p= 0.2126; 11d: p >0.9999; 13d: p=0.9338, Fig.S4B) and exhibited similar motor performance(Fig.5A, B) before any experimental procedures. This step ruled out potential confounding effects of the ABX on the general condition of the animals. Interestingly, NLRP3 inflammasome activation in the colon was suppressed when the gut microbiota after ICH was depleted, as shown by decreased protein levels of NLRP3, cleaved Caspase-1 p20 and IL-1β (NLRP3: Mann-Whitney U =0, p = 0.0286; Caspase-1 p20 : Mann-Whitney U = 0, p = 0.0286; IL-1β: t_6_ = 3.711, p=0.01, Fig.4B). This was accompanied by the preservation of the intestinal barrier, indicated by a significant increase in Occludin levels (ZO-1: t_6_=1.174, p = 0.2849; Occludin : t_6_ = 3.053, p = 0.0224, Fig. 4C). Transcriptional analysis corroborated these findings, showing downregulation of *Nlrp3* and *Asc* and upregulation of *Tjp1* and *Ocln* in colon tissue (*Nlrp3*: Mann-Whitney U =0, p = 0.0286; *Casp1*: Welch-corrected t_3.066_ = 2.474, p = 0.0879; *Asc* : t_5_ =2.735, p = 0.041; *Tjp1*: t_10_ = 3.055, p = 0.0122; *Ocln*: t_10_ =2.842, p = 0.0175, Fig.S4F-G).

**Fig 5.**
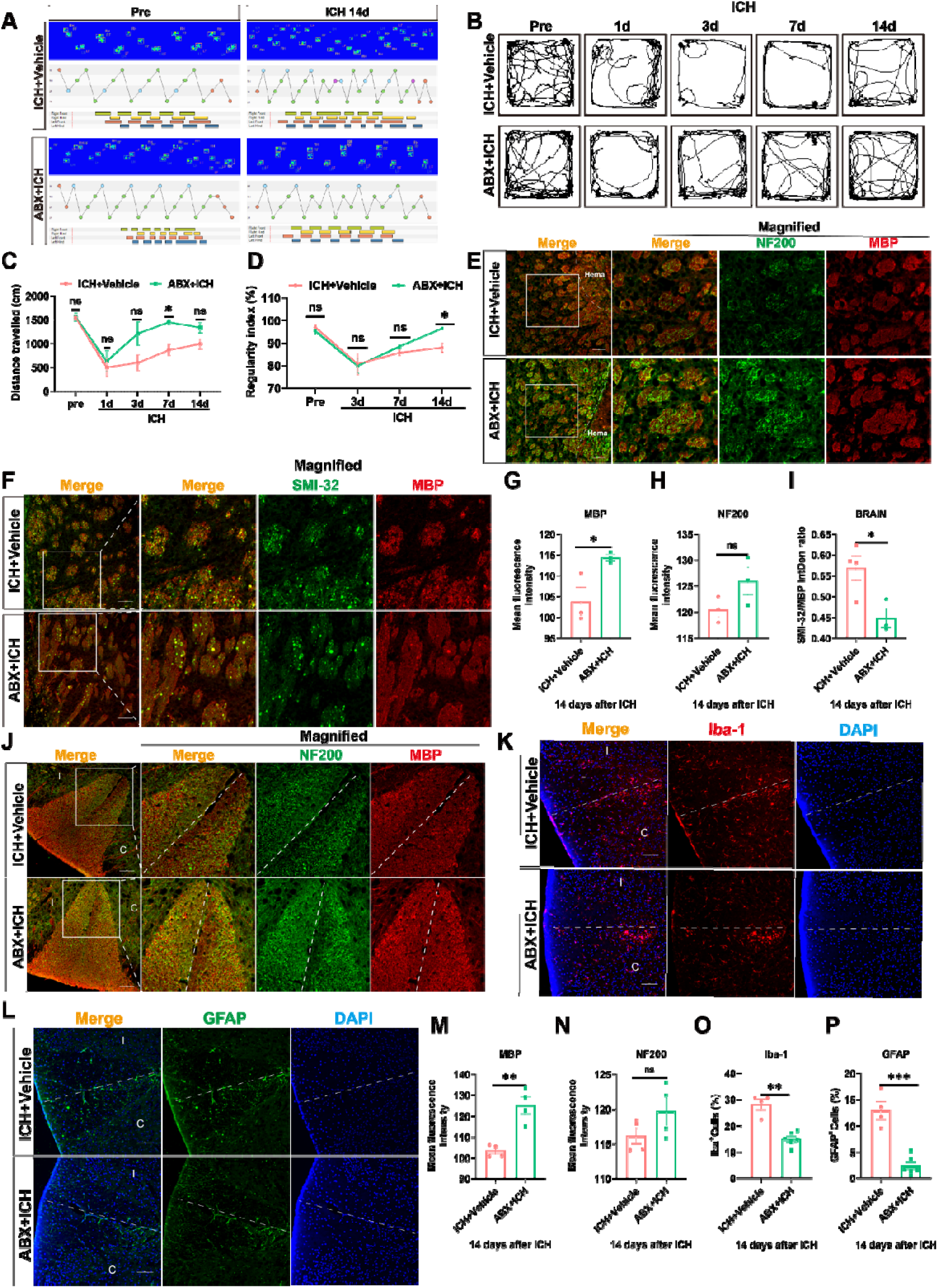
Gut microbiota depletion mitigated neurological deficits and CST injury after ICH. **(A, C)** Representative movement traces, and quantification of total distance traveled at 1d, 3d, 7d, and 14 days post-ICH (n=6 mice/group). Statistical analysis was performed using two-way ANOVA and Sidak’s multiple comparisons test. **(B, D)** Representative paw-print sequences and quantification of the Regularity Index at 3d, 7d, and 14 days post-ICH (n=5 mice/group). Statistical analysis was performed using two-way ANOVA and Sidak’s multiple comparisons test. (**E, G, H**) Representative immunofluorescence images and quantification of MBP and NF200 in the peri-hematoma brain region (n=3 mice/group). Scale bar: 100 μm. Statistical analysis by the two-tailed t test. (**F, I**) Representative immunofluorescence staining and quantification for MBP and SMI-32 in the peri-hematoma area (n=3-4 mice/group). Scale bar: 100 μm. Statistical analysis by the two-tailed t test. **(J, N, O)** Representative immunofluorescence images and quantification of MBP and NF200 in the cervical spinal cord (n=4 mice/group). Scale bar: 100 μm. An unpaired t test was used for MBP and NF200. Statistical analysis by the two-tailed t test. **(K, L, O, P)** Representative immunofluorescence images and quantification of Iba-1 (microglia) and GFAP (astrocytes) in the cervical spinal cord (n=4-6 mice/group). Scale bar: 100 μm. The Mann-Whitney test was used for Iba-1, while the unpaired t test was used for GFAP. Data are presented as mean ± SEM. Significance levels are defined as *p < 0.05, **p < 0.01, ***p < 0.001, and ****p < 0.0001.

The anti-inflammatory effects of microbiota depletion were systemic. In the peripheral blood, ABX treatment significantly lowered serum concentrations of IL-1β and Caspase-1 (Fig. 4D-E). Furthermore, in the ipsilateral brain hemisphere, ABX suppressed the NLRP3 inflammasome pathway at both the protein (NLRP3: t_6_=6.136, p = 0.0009; Caspase-1 p20 : t_6_= 4.268, p = 0.0053; IL-1β: t_6_ = 2.614, p=0.0399, Fig.4D-E) and mRNA levels (*Nlrp3*: t_4_=6.051, p = 0.038; *Casp1*: t_4_ = 3.672, p = 0.0214; *Asc* : t_6_ =3.120, p = 0.0206; *IL-1*β: t_5_ =3.062, p = 0.028, Fig.S4D), and ameliorated BBB disruption by increasing ZO-1 expression (ZO-1: t_6_=2.740, p = 0.0337; Occludin : t_6_= 2.257, p = 0.0648; *Tjp1*: t_5_ = 5.214, p = 0.0034; *Ocln*: Welch-corrected t_2.076_ =4.405, p = 0.0447, Fig.4F-G and Fig.S4E).

To summarize, these results establish that the gut microbiota is essential for driving NLRP3 inflammasome activation after ICH. Depleting the gut microbiota after ICH interrupted this pathway at its origin in the gut, leading to attenuated inflammatory signaling in the circulation and the brain. This cascade logically positions the microbiota as an upstream regulator of the CST injury and functional deficits observed after ICH.

### 3.6. Gut microbiota depletion mitigates neurological deficits and CST injury after ICH

We next sought to determine the functional consequences of gut microbiota depletion on neurological outcomes and CST integrity. Behavioral assessments revealed that ABX treatment significantly improved motor function after ICH. ABX-ICH group exhibited increased total distance traveled in the open-field test at 3 days post-ICH (pre: p>0.9999; 1d: p=0.9956; 3d: p=0.3333; 7d: p=0.0173; 14d: p=0.2488, Fig.5A, C) and showed a higher regularity index in CatWalk XT™ gait analysis, indicating improved neurological function (pre: p=0.8224; 3d: p=0.9999; 7d: p=0.3769; 14d: p=0.0421, Fig.5B, D).

We then assessed the structural correlates of this functional improvement. In the peri-hematomal region, ABX treatment significantly attenuated ICH-induced demyelination, as shown by preserved MBP expression (MBP: t_4_=3.018, p=0.0392; NF200: t_4_=1.804, p=0.1455, Fig.5E, G, H). Furthermore, SMI-32/MBP co-staining confirmed that ABX significantly reduced the proportion of degenerating axons within the CST (SMI-32/MBP: t_5_=3.067, p=0.0279, Fig.5F, I), despite a non-significant trend in NF200 expression (Fig.5H). Besides, ABX treatment mitigated the loss of MBP (MBP: t_6_=5.160, p=0.0021; NF200: t_6_=1.574, p=0.1666, Fig.5J, M, N) of the cervical spinal cord and suppressed neuroinflammation, as evidenced by reduced activation of Iba-1□ microglia and GFAP□ astrocytes (Iba-1: Mann-Whitney U=0, p=0.0095; GFAP: t_8_=6.752, p=0.0001, Fig.5K, L, O, P).

In summary, gut microbiota depletion confers comprehensive neuroprotection, improving motor recovery and preserving CST integrity both at the primary injury site and in the distal spinal cord. These findings definitively establish the gut microbiota as a critical upstream driver of post-ICH pathology and functionally link its role to the previously observed gut NLRP3 inflammasome-dependent pathway of CST injury.

### 3.7. Gut NLRP3 activation is dependent on gut microbiota dysbiosis to exacerbate CST injury after ICH

Although our data implicated both gut dysbiosis and gut NLRP3 activation in post-ICH pathology, their causal interplay remained undefined. To directly establish the causal role of the gut NLRP3 inflammasome in microbiota-mediated exacerbation of CST injury, we employed a combined genetic and microbiological intervention (Fig.6A). Importantly, the groups showed comparable baseline body weight and motor function before ICH induction (Fig.S5A and Fig.6D, E), indicating that the differential neurological outcomes were not due to differences in health status before ICH. After the gut microbiota was cleared by antibiotics, colonic-specific NLRP3 knockdown (AAV-NLRP3) or control (AAV-GFP) mice received fecal microbiota transplantation (FMT) from ICH donors. Among them, the AAV-GFP control group was further divided into two groups: one received FMT from ICH donors (AAV-GFP_FMT-ICH) to control the microbial composition, and the other only received the vehicle (AAV-GFP_Vehicle) to establish a baseline state of microbiota depletion. This design aimed to investigate whether knockdown of intestinal NLRP3 could reverse the harmful effects of the ICH microbiota.

**Fig 6.**
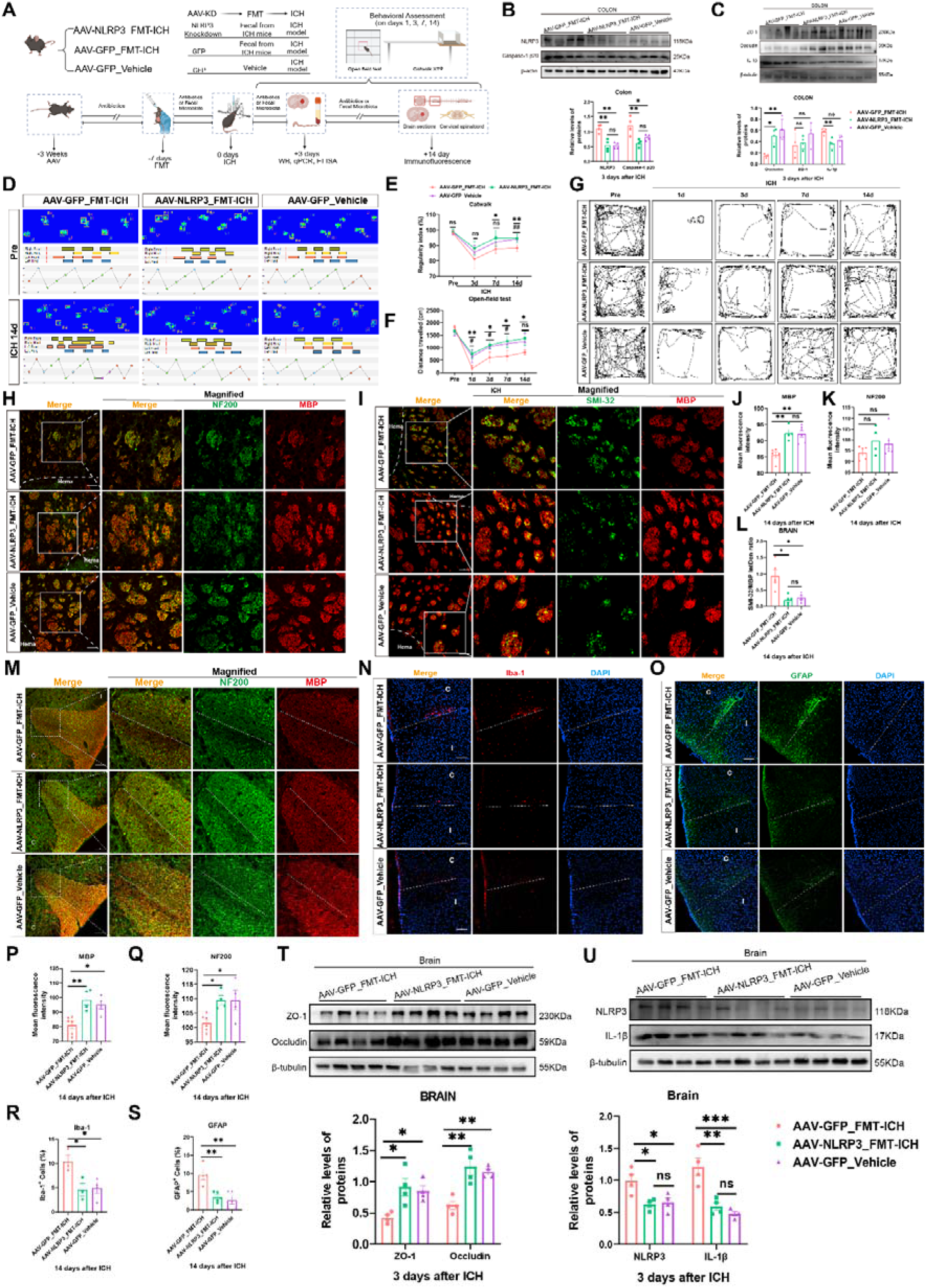
Gut NLRP3 activation was dependent on gut microbiota dysbiosis to exacerbate CST injury after ICH. **(A)** Schematic of experimental design. **(B-C)** Colon NLRP3 inflammasome components (NLRP3, Caspase-1 p20, IL-1β) and gut barrier (ZO-1 and Occludin) were analyzed by Western blot. n=4 mice/group. Statistical analysis was performed using the one-way ANOVA test followed by Tukey’s post hoc test for multiple comparisons. **(D-E)** Representative footprints and the regularity Index from the CatWalk XT analysis, n=6 mice/group. Statistical analysis was performed using the two-way ANOVA test with Tukey’s multiple comparisons test. **(F-G)** Representative movement traces and quantification of total distance traveled at 1d, 3d, 7d, and 14 days post-ICH. n=4-5 mice/group. Statistical analysis was performed using the two-way ANOVA test followed by Tukey’s post hoc test for multiple comparisons. **(H, J, K)** MBP and NF200 immunofluorescence staining in peri-hematoma brain. n=4-6 mice/group. Scale bar:100 μm. One-way ANOVA test with Tukey’s multiple comparisons test was used for MBP, while Kruskal-Wallis test with Dunn’s multiple comparisons test was used for NF200. (**I, L**) MBP and SMI-32 immunofluorescence in peri-hematoma brain. n=5mice/group. Scale bar:100 μm. Statistical analysis by Welch’s ANOVA test with Games-Howell’s multiple comparisons test. **(M, P, Q)** MBP and NF200 immunofluorescence in the cervical spinal cord (n=4-6 mice/group). Scale bar: 100 μm. One-way ANOVA with Tukey’s multiple comparisons test was used for MBP and NF200. **(N, O, R, S)** Representative Iba-1 and GFAP immunofluorescence staining were performed on cervical spinal cord sections. n=3-5 mice/group. Scale bar:100 μm. One-way ANOVA with Tukey’s multiple comparisons test was used for Iba-1 and GFAP. **(T-U)** Western blot analysis of brain NLRP3, IL-1β, ZO-1 and Occludin. n=4 mice/group. One-way ANOVA with Tukey’s multiple comparisons test was used for these. β-tubulin or β-actin was used as the loading control in western blot analysis. All data are presented as meanLJ±LJSEM. *: AAV-GFP_FMT-ICH vs. AAV-NLRP3_FMT-ICH; #: AAV-GFP_FMT-ICH vs. AAV-GFP_Vehicle; *, ^#^p < 0.05, **, ^##^p < 0.01, ***, ^###^p < 0.001.

**Fig 7.**
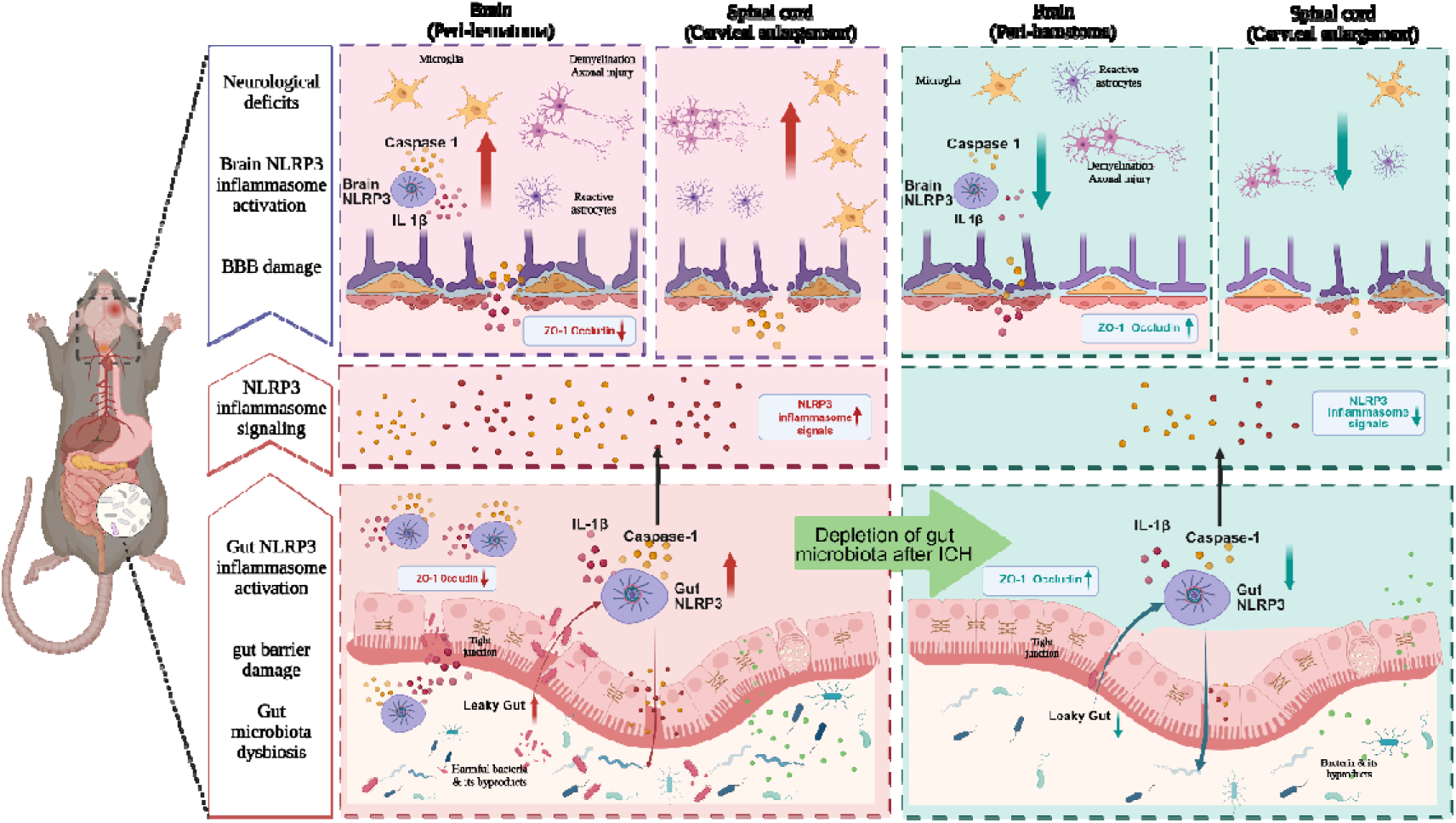
Gut Microbiota Dysbiosis-Mediated Gut NLRP3 Inflammasome Activation Exacerbates Corticospinal Tract Injury After Intracerebral Hemorrhage. Schematic depicting a gut-brain pathway that exacerbates CST injury after ICH. ICH initially causes gut microbiota dysbiosis, which activates the gut NLRP3 inflammasome and impairs the gut barrier. The resulting inflammatory signals lead to the systemic dissemination of inflammatory mediators. These signals subsequently aggravate BBB disruption and peri-hematomal NLRP3 activation, ultimately amplifying CST injury. The figure was created with BioRender (https://BioRender.com/4wz3lug).

The results indicated that mice receiving the ICH microbiota but with intact intestinal NLRP3 (AAV-GFP_FMT-ICH) exhibited significant colon inflammasome activation, with protein levels of NLRP3, cleaved Caspase-1 p20 and IL-1β all significantly higher than those in the baseline control group (AAV-GFP_Vehicle) (NLRP3: F_(2, 9)_ = 19.31, p=0.0006; Caspase-1 p20: F_(2, 9)_ = 9.429, p= 0.0062; IL-1β: F _(2, 9)_ = 7.913, p=0.0104, Fig. 6B, C). Crucially, colon NLRP3 knockdown (AAV-NLRP3_FMT-ICH group) blocked this activation, reducing the above inflammatory markers to levels indistinguishable from the baseline control group that did not receive the ICH microbiota (Fig. 6B, C). Tight junction protein Occludin also showed a similar protective pattern, while ZO-1 did not change significantly among the groups (Occludin: F _(2, 9)_ = 13.26, p=0.0021; ZO-1: F _(2, 9)_ = 1.452, p=0.2841, Fig. 6C).

Subsequently, we confirmed that the colon NLRP3 knockdown can effectively alleviate CST damage. In terms of neurological function, the AAV-GFP_FMT-ICH group showed the most severe impairment, characterized by a decrease in the regularity index of motor function (7d: AAV-GFP_FMT-ICH vs. AAV-NLRP3_FMT-ICH, p= 0.0477; 14d: AAV-GFP_FMT-ICH vs. AAV-NLRP3_FMT-ICH, p= 0.0046; AAV-GFP_FMT-ICH vs. AAV-GFP_Vehicle: p=0.0048. Fig. 6D, E) and a reduction in the total movement distance in the open-field (1d: AAV-GFP_FMT-ICH vs. AAV-NLRP3_FMT-ICH, p= 0.0025; AAV-GFP_FMT-ICH vs. AAV-GFP_Vehicle: p=0.0253. 3d: AAV-GFP_FMT-ICH vs. AAV-NLRP3_FMT-ICH, p= 0.0267; AAV-GFP_FMT-ICH vs. AAV-GFP_Vehicle: p=0.045. 7d: AAV-GFP_FMT-ICH vs. AAV-NLRP3_FMT-ICH, p= 0.0175; AAV-GFP_FMT-ICH vs. AAV-GFP_Vehicle: p=0.0315. 14d: AAV-GFP_FMT-ICH vs. AAV-NLRP3_FMT-ICH, p= 0.0198. Fig. 6F, G). Both knockdown of gut NLRP3 and simple removal of the microbiota provided significant and comparable protection, enabling the mice to perform better than the AAV-GFP_FMT-ICH group (Fig. 6D-G).

Histological analysis revealed that the AAV-GFP_FMT-ICH group showed the most severe injury to the CST surrounding the hematoma, manifested by significant demyelination and axonal degeneration (MBP: F _(2, 12)_ = 13.54, p=0.0008; NF200: Kruskal-Wallis H=1.028, p=0.6216; SMI-32/MBP: F _(2, 7.316)_ = 6.516, p=0.0086, Fig. 6H-L). Besides, the AAV-GFP_FMT-ICH group had significantly lower levels of both MBP and NF200 compared to the AAV-NLRP3_FMT-ICH and AAV-GFP_Vehicle groups in cervical CST (MBP: F _(2, 11)_ = 10.39, p=0.0029; NF200: F _(2, 11)_ =5.661, p=0.0204, Fig. 6M, P, Q). Similarly, both protective interventions were effective in preventing the above structural damage, and there was no difference in the effects of the two (Fig. 6H-M, P, Q).

Ultimately, this protective cascade reaction significantly alleviated neuroinflammation in cervical CST and maintained the integrity of the BBB. The AAV-GFP_FMT-ICH group showed significantly higher activation of microglia (Iba-1) and astrocytes (GFAP) in the cervical spinal cord (Iba-1: F (2, 7) = 6.564, p=0.0248; GFAP: F (2, 11) = 12.85, p=0.0013. Fig. 6N, O, R, S), while the two protective groups effectively inhibited this reaction. In the brain, the harmful effects of the ICH microbiota were associated with increased disruption of the BBB (decreased ZO-1 and Occludin) and activation of the NLRP3 inflammasome in the brain, and knocking down intestinal NLRP3 or eliminating the ICH microbiota could significantly alleviate these pathological changes (ZO-1: F _(2, 9)_ = 7.305, p=0.013; Occludin: F _(2, 9)_ = 14.18, p=0.0017; NLRP3: F _(2, 9)_ = 7.833, p=0.0107. IL-1β: F _(2, 9)_ = 19.54, p=0.0005. Fig.6T-U).

Taken together, these data provide definitive evidence that ICH-related microbiota activate the gut NLRP3 inflammasome, which in turn exacerbates gut barrier disruption, neuroinflammation, and CST injury. The finding that preventing gut microbiota dysbiosis also reaches the protective effect of gut NLRP3 knockdown strongly suggests that the detrimental role of gut NLRP3 in ICH pathology may depend on gut microbiota dysbiosis.

## 4. Discussion

ICH leads to severe and persistent neurological deficits, with secondary CST injury being a critical determinant of long-term functional outcome ^[2]^. Although previous studies mainly focused on the pathological changes around the hematoma, our research team previously emphasized that distal cervical enlargement spinal cord injury might be a key factor leading to persistent injury ^[3b, 16, 18]^. However, the mechanisms remain poorly defined.

The gut-brain axis has emerged as a key regulator of neurological health and disease ^[19]^. Here, we identify a specific gut-brain pathway that significantly worsens CST injury after ICH. We have found that ICH can cause certain changes in the gut, including alterations in the gut microbiota, activation of the NLRP3 inflammasome in the gut, and damage to the intestinal barrier. By knocking down the NLRP3 gene in the colon or using antibiotics to eliminate the intestinal microbiota, we established a clear causal chain. The imbalance of gut microbiota activates the NLRP3 inflammasome in the colon. This mechanism can damage the intestinal barrier, allowing inflammatory mediators such as Caspase-1 and IL-1β to enter the peripheral blood. These factors will further damage the BBB, intensify the inflammatory response around the brain injury, and ultimately lead to the degeneration of axons along with the loss of myelin.

Contrary to the traditional view that higher alpha diversity indicates a healthier biological system, we observed a significant increase in multiple alpha diversity indices (Richness, Chao1, ACE, and Shannon) following ICH. Although this phenomenon may seem contrary to common sense, it is increasingly being recognized in other acute inflammatory diseases as well; it can be understood as a sign of severe imbalance in the microbial ecosystem ^[20]^. We propose that the ICH-induced disruption of the gut environment leads to a collapse of dominant commensal populations, thereby permitting the uncontrolled expansion of opportunistic taxa (e.g., *Akkermansia*, *Prevotella*). Consequently, the observed increase in alpha diversity does not reflect a healthier state but rather a bloom of inflammation-associated opportunistic taxa, which coincides with a loss of commensal bacteria (e.g., *Muribaculum*, *Paramuribaculum*). Several recent studies focusing on ICH have consistently reported a significant increase in the abundance of *Akkermansia*, which aligns with our own results ^[5, 21]^. This bacterium uses mucin O-glycans as its main carbon source, and these mucin O-glycans are the main components of the intestinal mucus layer. Moreover, this bacterium also contains abundant sialidase ^[22]^. Excessive degradation of mucins by the microbiota can disrupt the gut barrier and thereby exacerbate intestinal inflammation ^[21a]^. In contrast, the abundance of *Paramuribaculum* and *Muribaculum*, which are commensal bacteria that contribute to gut microbiota homeostasis, was reduced after ICH ^[23]^. This restructured community is likely more pro-inflammatory and contributes to the pathogenesis, as supported by our findings of gut NLRP3 inflammasome activation and barrier disruption.

One interesting finding in our research is that the NLRP3 inflammasome in the gut does not act independently. Its destructive effects may depend on the signals released by the gut microbiota dysbiosis after ICH. We reached this conclusion based on two key pieces of evidence. First, antibiotic-mediated depletion of the gut microbiota substantially decreased the activation of the colon NLRP3 inflammasome post-ICH. More importantly, when we transplanted the disturbed microbiota from ICH mice into recipient mice, it caused severe pathology only if the recipients’ gut NLRP3 gene was intact. If the recipients lacked gut NLRP3, they were protected despite receiving the same microbiota from ICH. This discovery demonstrates the crucial role of the gut NLRP3 inflammasome in the process by which gut microbiota dysbiosis causes CST damage after ICH, and it is a necessary mediator factor in this pathological process.

Our findings significantly advance the field by moving beyond correlative observations to establish a causal gut-brain axis in ICH pathology. Although our previous studies on MCC950 and oxymatrine had already suggested a certain connection between NLRP3 inhibition, changes in the microbiota, and neuroprotective effects, this current study finally fills in this gap in the mechanism ^[3b]^. From the translational perspective, our data suggest that interventions targeting the gut microenvironment, either by resolving dysbiosis or by locally inhibiting the gut NLRP3 inflammasome could offer a strategic and potentially safer avenue for mitigating CST injury after ICH.

While this study systematically elucidates the gut microbiota-NLRP3 axis in CST injury post-ICH, several limitations warrant consideration. Firstly, relying on young adult mice as the experimental model may limit the application of these research results in clinical practice. This is not only because there are differences in the composition of gut microbiota, immune responses, and BBB structure among different species, but also because the age of the experimental animals does not match the elderly population in which ICH cases mainly occur in clinical practice. Future research can use elderly mouse models and collect human samples for further verification, which will help overcome this limitation. Secondly, although ABX depletion validated the functional role of gut microbiota, it failed to identify specific bacterial taxa. We will explore this in our next work. Finally, our data indicate that the activation of the systemic inflammasome is associated with pathological changes in the central nervous system. It is still necessary to verify whether a significant portion of the Caspase-1/IL-1β substances present in the circulatory system actually originates from the gut. This issue remains to be confirmed. The known phenomenon of immune cells from the gut to the brain after ICH offers a potential mechanism worthy of further investigation ^[5, 24]^. We will conduct further exploration and analysis in our subsequent research.

In summary, we have delineated a complete gut-brain axis that exacerbates CST injury after ICH. This pathway begins with gut microbiota dysbiosis, which subsequently triggers the activation of the NLRP3 inflammasome in the intestine, ultimately leading to damage to the distal spinal cord. By mapping this route, our work not only deepens the understanding of the pathological mechanism of the CST after ICH, but also points out the direction for new intervention measures aimed at promoting brain recovery by regulating gut function. Future studies aimed at identifying the specific bacterial taxa and metabolites responsible for NLRP3 activation, as well as directly tracing the trafficking of gut-derived inflammatory mediators to the brain, will be crucial to fully exploit the therapeutic potential of this pathway.

## Data availability

All data are available upon reasonable request. The 16S rRNA sequencing data generated in this study are deposited in the National Genomics Data Center, China National Center for Bioinformation/Beijing Institute of Genomics, Chinese Academy of Sciences (GSA: CRA034111).

## Acknowledgements

We thank Dr. Xiaolong He for helpful discussions and insightful comments. The image of the mechanism was created with BioRender.com. During the preparation of this work, the authors used DeepSeek to improve readability and language fluency. After using this tool, the authors reviewed and edited the content as needed and take full responsibility for the content of the publication.

## Author contributions

H.S.: Conceptualization, Supervision, Project Administration, Funding Acquisition, Writing – Review & Editing. J.H.: Supervision, Project Administration, Writing – Review & Editing. M.Z. and M.P.: Investigation, Formal Analysis, Writing – Original Draft. H.T., C.H., and L.Z.: Investigation (specifically in histology, qPCR, behavior tests, and Western blot). W.Y.: Resources, Investigation (behavior tests). All authors reviewed and approved the final manuscript.

