## Supplementary Figures and table for "Gut Microbiota Dysbiosis-Mediated Gut NLRP3 Inflammasome Activation Exacerbates Corticospinal Tract Injury After Intracerebral Hemorrhage"

**Supplementary figures and figure legends：**

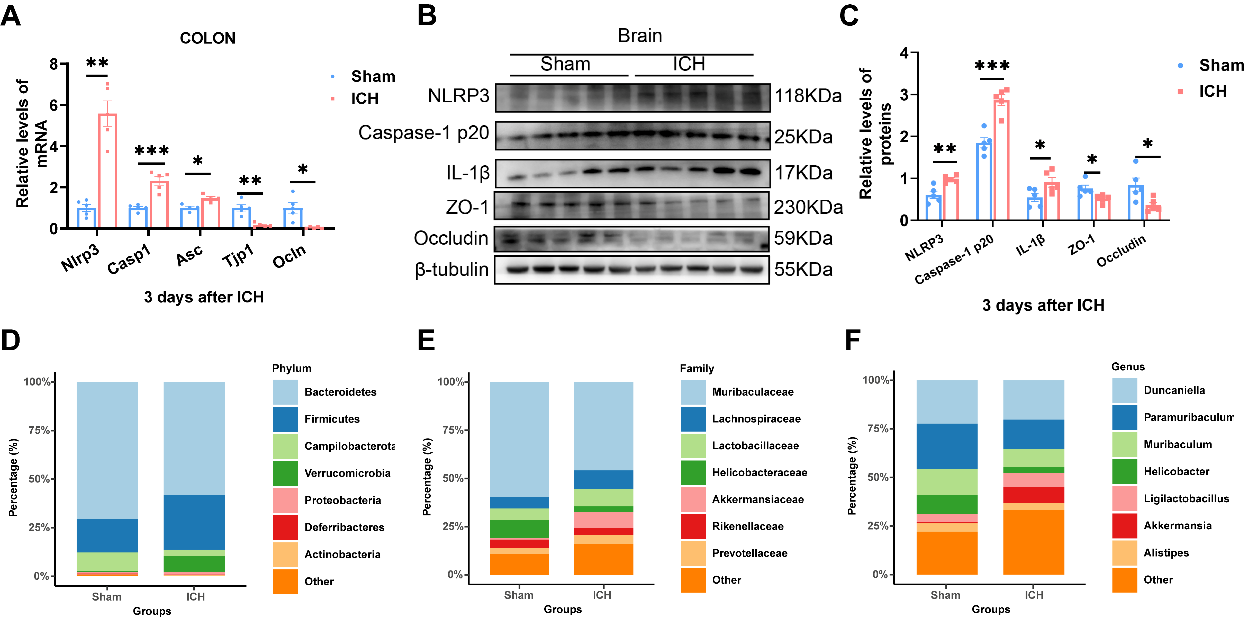

**Supplementary Fig. 1 Pathological features of the gut-brain axis post-ICH, including microbiota dysbiosis, NLRP3 activation, and barrier disruption. (A)** The mRNA expression level of *Nlrp3*, *Casp1*, *Asc,* *Tjp1* and *Ocln* in colon (n=4-5 mice/group). Statistical analysis by the two-tailed t test (*Casp1* and *Asc*) and two-tailed t test with welch’s correction

(*Nlrp3, Tjp1* and *Ocln*). **(B-C)** Western blots of brain NLRP3 inflammation proteins (NLRP3, Caspase-1 p20, IL-1β) and tight junction proteins (Occludin, ZO-1). β-tubulin was used as a loading control, n=5 mice/group. Statistical analysis by the two-tailed t test. **(D-F)** Compositional profile of the gut microbiota shown by stacked bar plots, depicting the relative abundance of the top seven most abundant taxa at the phylum (D), family (E), and genus (F) levels, n=8-9 mice/group. All data were analyzed using the unpaired t test. Data were considered significant if *p < 0.05, **p < 0.01, ***p < 0.001, ns no significance.

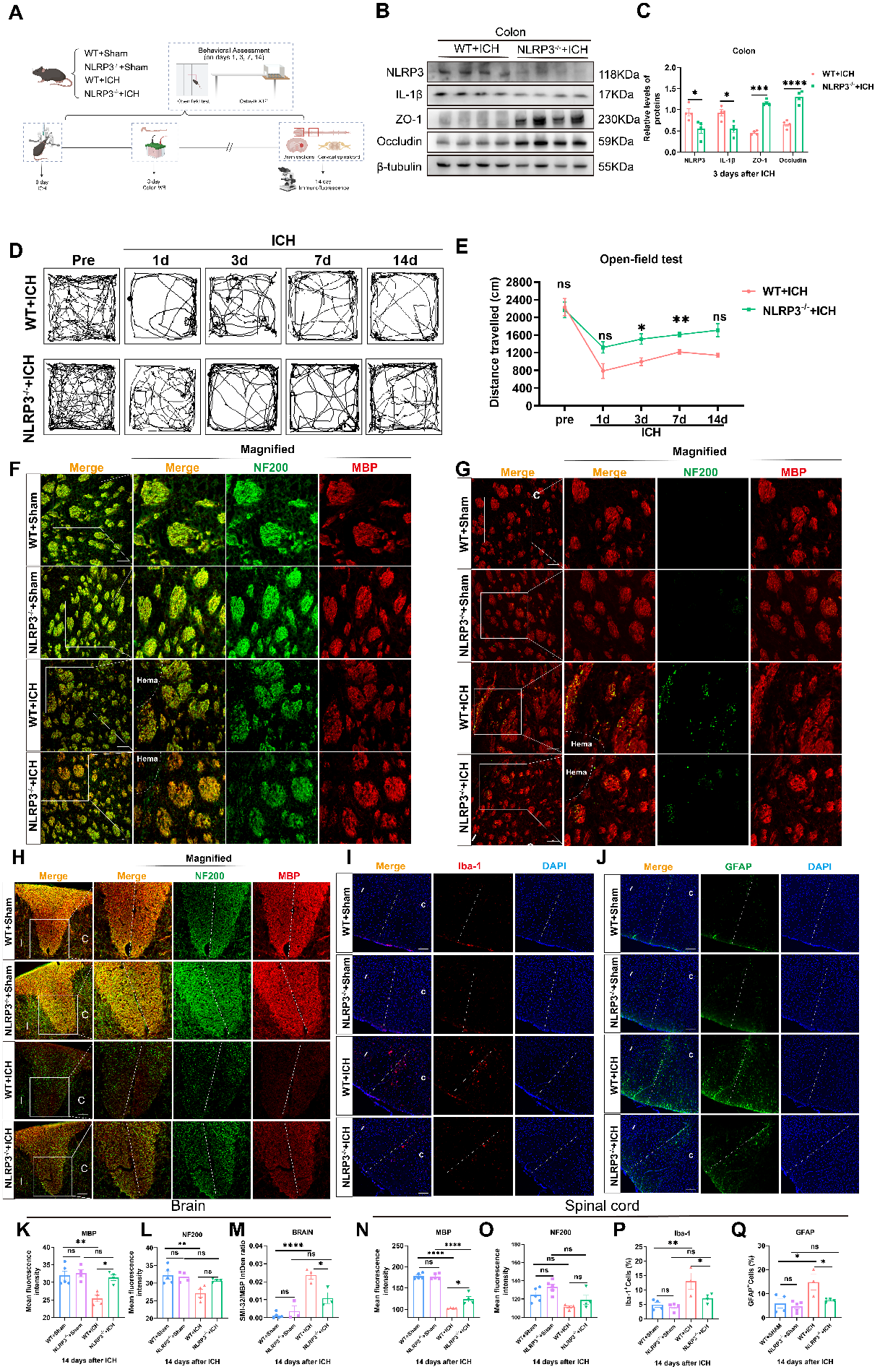

**Supplementary Fig. 2 NLRP3 knockout ameliorates brain hematoma volume and mitigates CST injury after ICH.** **(A)** Experimental schematic outlining the timeline, treatment groups, and procedures (https://BioRender.com/3bls52h). **(B-C)** Western blots of brain colon inflammation proteins (NLRP3, IL-1β) and tight junction proteins (Occludin, ZO-1). β-tubulin was used as the loading control, n=4/group. Statistical analysis by the two-tailed t test. **(D, E)** Open field test: (D) Representative locomotion trajectories and (E) quantification of total distance traveled over 14 days post-ICH (n=6 /group). Statistical analysis by the two-way ANOVA with Sidak's multiple comparisons test. **(F, K-L)** Representative immunofluorescence images and quantification of MBP and NF200 in the peri-hematoma brain region (n=4-5 mice/group). Scale bar, 100 μm. **(G, M)** Representative immunofluorescence images and quantification of SMI-32 and MBP in the cervical spinal cord (n=3-5 mice/group). Scale bar, 100 μm. **(H, N-O)** Representative MBP and NF200 immunofluorescence staining in the cervical spinal cord. n=4-5 mice/group. Scale bar:100 μm. **(I-J, P-Q)** Representative immunofluorescence images and quantification of Iba-1 and GFAP in the cervical spinal cord (n=3-5 mice/group). Scale bar, 100 μm. Data are presented as mean ± SEM. Statistical analysis was performed using one-way ANOVA with Tukey's multiple comparisons test for K-Q. *p < 0.05, **p < 0.01, ***p < 0.001, ****p < 0.0001. C: contralateral; I: ipsilateral, ns: no significance.

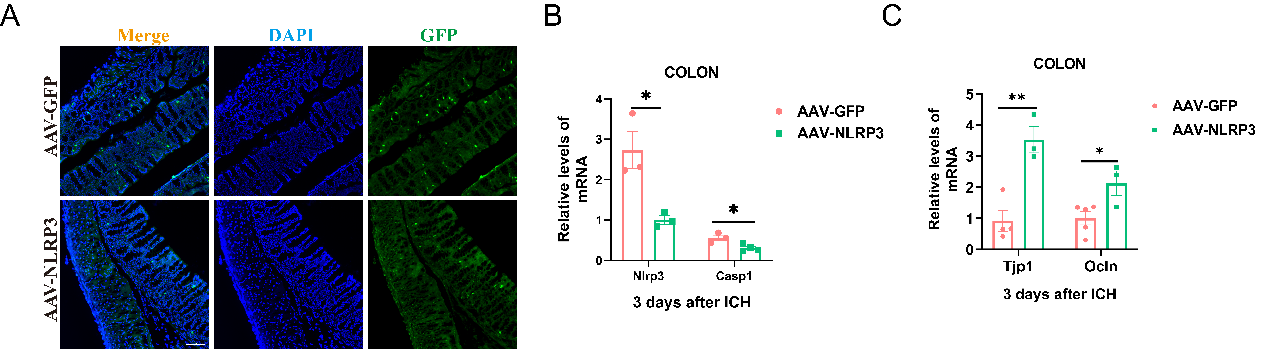

**Supplementary Fig. 3 Validation of colon-specific NLRP3 knockdown and its functional impact. (A)** Immunofluorescence image validating successful AAV transduction in colon tissue**. (B)** qPCR analysis confirms efficient knockdown of *Nlrp3* and the concomitant downregulation of *Casp1* mRNA expression in the colon, n=3-4/group**.** Statistical analysis by the two-tailed t test. **(C)** NLRP3 knockdown alleviated ICH-induced damage to the intestinal barrier (*Tjp1*, *Ocln*), n=3-5/group. Statistical analysis by the two-tailed t test. Data were analyzed using the unpaired t test. Data were considered significant if *p < 0.05, **p < 0.01, ***p < 0.001, ****p < 0.0001, ns no significance.

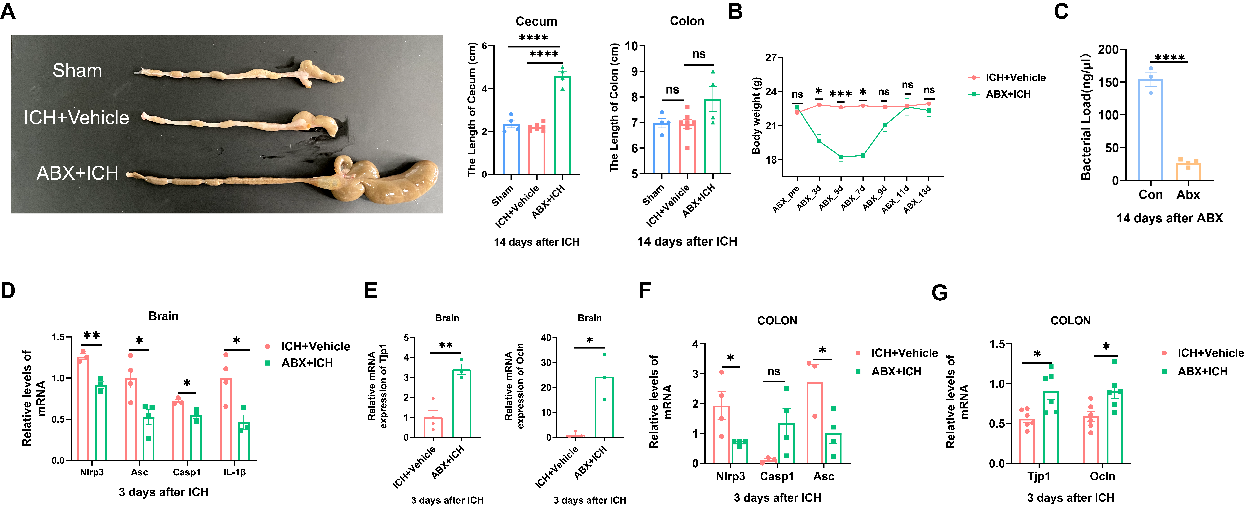

**Supplementary Fig. 4 Gut Microbiota Depletion Alters NLRP3 Inflammasome in the Gut and Brain. (A)** ABX treatment induced significant cecal enlargement (n=4-7 mice/group). Statistical analysis was performed using the one-way ANOVA and Tukey's multiple comparisons test. **(B)** Body weight changes in mice after ABX administration (n=6 mice/group). Statistical analysis was performed using the two-way ANOVA test and Sidak's multiple comparisons test. **(C)** ABX treatment markedly reduced the gut bacterial load (n=3-4 mice/group). Statistical analysis used the two-tailed t test. **(D-E)** ABX treatment downregulated the mRNA expression of NLRP3 inflammasome components (*Nlrp3*, *Asc*, *Casp1*, *Il-1β*) and tight junction proteins (*Tjp1*, *Ocln*) in the brain (n=3-4 mice/group). Statistical analysis by the two-tailed t test in other panels, t test with Welch's correction in *Ocln*. **(F-G)** ABX treatment reduced the mRNA levels of *Nlrp3*, *Asc*, *Casp1*, *Tjp1*, and *Ocln* in the colon (n=4-5 mice/group). Data are presented as mean ± SEM. Mann-Whitney test was used for *Nlrp3*, t test with Welch's correction was used for *Casp1* and two-tailed t test was used for *Asc*, *Tjp1* and *Ocln*. Significance was set at *p < 0.05, **p < 0.01, ***p < 0.001, and ****p < 0.0001, ns no significance.

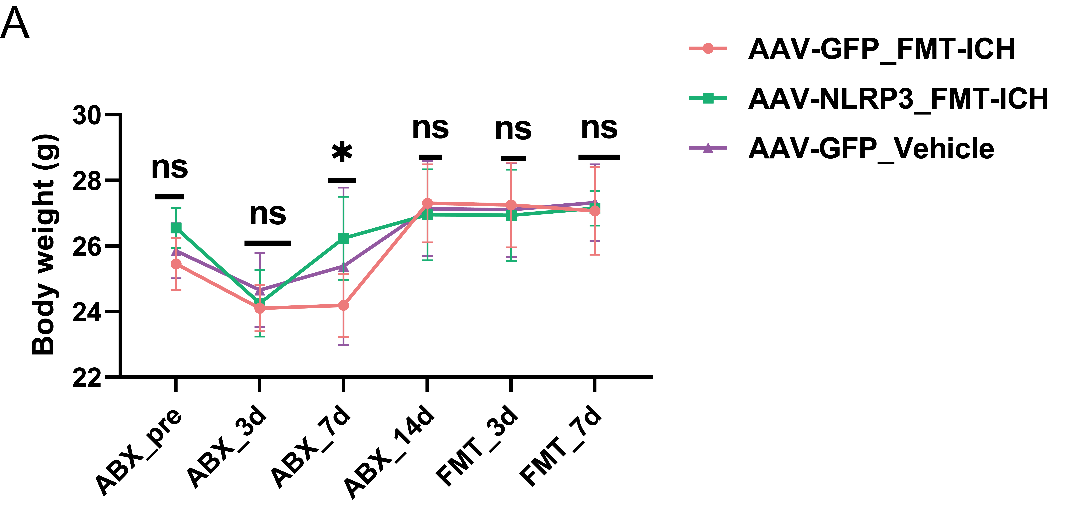

**Supplementary Fig. 5 Baseline body weight before ICH induction. (A)** Body weight changes in mice after ABX and FMT administration (n=6 mice/group). Statistical analysis was performed using the two-way ANOVA test and Tukey's multiple comparisons test. *p < 0.05, **p < 0.01, ***p < 0.001, ****p < 0.0001, ns no significance.

**Supplementary table：**

Supplementary table 1. Primers used in this study.

| Organism | Target | Sequence (5’-3’) |
| --- | --- | --- |
| *Mus musculus* | *Nlrp3* | F：CCCCGCCATGTGGAGATCCTAGG |
|  |  | R：GGCCAGGCTCTTCCCGGTCTCC |
| *Mus musculus* | *Casp1* | F：CTTGGAGACATCCTGTCAGGG |
|  |  | R：AGTCACAAGACCAGGCATATTCT |
| *Mus musculus* | *Asc* | F：GACAGTGCAACTGCGAGAAG |
|  |  | R：CGACTCCAGATAGTAGCTGACAA |
| *Mus musculus* | *Il-1b* | F：AAGGGCTGCTTCCAAACCTTTGAC |
|  |  | R：TGCCTGAAGCTCTTGTTGATGTGC |
| *Mus musculus* | *Tjp1* | F：GGGGCCTACACTGATCAAGA |
|  |  | R：TGGAGATGAGGCTTCTGCTT |
| *Mus musculus* | *Ocln* | F：ACGGACCCTGACCACTATGA |
|  |  | R：TCAGCAGCAGCCATGTACTC |
| *Mus musculus* | *Actb* | F：AGAGGGAAATCGTGCGTGAC |
|  |  | R：CAATAGTGATGACCTGGCCGT |

*Nlrp3*, NOD-like receptor thermal protein domain-associated protein 3*; Casp1,* Caspase-1; *Asc*, apoptosis-associated speck-like protein containing a CARD; *Il-1b*, interleukin 1 beta; *Tjp1*, Tight junction protein 1; *Ocln*, Occludin; *Actb*, actin beta.
